# Mitochondrial protein import clogging as a mechanism of disease

**DOI:** 10.1101/2022.09.20.508789

**Authors:** Liam P. Coyne, Xiaowen Wang, Jiyao Song, Ebbing de Jong, Karin Schneider, Paul T. Massa, Frank A. Middleton, Thomas Becker, Xin Jie Chen

## Abstract

Mitochondrial biogenesis requires the import of >1,000 mitochondrial preproteins from the cytosol. Most studies on mitochondrial protein import are focused on the core import machinery. Whether and how the biophysical properties of substrate preproteins affect overall import efficiency is underexplored. Here, we show that protein traffic into mitochondria is disrupted by amino acid substitutions in a single substrate preprotein. Pathogenic missense mutations in adenine nucleotide translocase 1 (Ant1), and its yeast ortholog Aac2, cause the protein to accumulate along the protein import pathway, thereby obstructing general protein translocation into mitochondria. This impairs mitochondrial respiration, cytosolic proteostasis and cell viability independent of Ant1’s nucleotide transport activity. The mutations act synergistically, as double mutant Aac2/Ant1 cause severe clogging primarily at the Translocase of the Outer Membrane (TOM) complex. This confers extreme toxicity in yeast. In mice, expression of a super-clogger Ant1 variant led to an age-dependent dominant myopathy that phenocopies Ant1-induced human disease, suggesting clogging as a mechanism of disease. We propose that secondary structures of mitochondrial preproteins play an essential role in preventing clogging and disease.

## Introduction

Mitochondria are essential organelles responsible for a wide range of cellular functions. To carry out these functions, they are equipped with a proteome of 1,000-1,500 proteins (1–4). The vast majority of these proteins are encoded by the nuclear genome, synthesized in the cytosol, and sorted into one of the four mitochondrial sub-compartments, namely the outer mitochondrial membrane (OMM), intermembrane space (IMS), inner mitochondrial membrane (IMM) and the matrix (5–8). The entry gate by which >90% of mitochondrial proteins enter mitochondria is the translocase of the outer membrane (TOM) complex. Therefore, proper function of the TOM complex is paramount for mitochondrial function and cell viability.

After passage through the TOM complex, specialized protein translocases transport preproteins into the mitochondrial sub-compartments (5–9). The import of mitochondrial carrier proteins to the protein-dense IMM is particularly challenging. Molecular chaperones like Hsp70 and Hsp90 target these highly hydrophobic proteins through the cytosol to the Tom70 receptor, which is associated with the TOM complex on the mitochondrial surface (10, 11). After transport through the channel-forming Tom40 subunit of the TOM complex, the heterohexameric small TIM chaperones (i.e. the Tim9-Tim10 complex) transport the preprotein to the carrier translocase of the inner membrane (TIM22 complex) (12–18). The carrier translocase is a multisubunit complex that inserts carrier proteins into the IMM in a membrane potential (Δψ)-dependent manner (19–24). The central subunit Tim22 integrates carrier proteins into the IMM (22, 25). It associates with different partner proteins in yeast and human mitochondria. In yeast, Tim18 and Sdh3 are required for assembly and stability of the carrier translocase, whereas Tim54 tethers the small TIM chaperones (Tim9-Tim10-Tim12 complex) to the translocase (26, 27). In human cells, TIM22 associates with the acylglycerol kinase (AGK) and TIM29, which are both required for full import capacity (28–31). Human TIM22 also associates with the TIM9-TIM10 complex (24).

Several diseases are associated with mutations directly affecting the protein import machinery of the carrier pathway. For example, mutations in *TIMM8A* have been found in patients suffering Mohr-Tranebjaerg syndrome/deafness dystonia syndrome (32). Mutations in *TOMM70* and *TIMM22* have been linked to a devastating neurological syndrome and a mitochondrial myopathy, respectively (33–35). Mutations in another TIM22 complex subunit, AGK, has been associated with the Senger’s syndrome marked by cataracts, hypertrophic cardiomyopathy, skeletal myopathy, and exercise intolerance (30, 31, 36). Clearly, defective protein import is linked to neurological and musculoskeletal diseases. However, whether protein import defects contribute to diseases not directly related to mutations in the core protein import machinery is unclear.

Here, we show that pathogenic missense mutations in a mitochondrial carrier protein, adenine nucleotide translocase 1 (Ant1) or ADP/ATP carrier 2 (Aac2) in yeast, cause arrest of the protein at the translocases during import into mitochondria. This effectively “clogs” the protein import pathway to obstruct general protein import and induce muscle and neurological disease in mice. Our findings demonstrate that global protein import is vulnerable to missense mutations in mitochondrial preproteins, and also provide strong evidence that protein import clogging contributes to neurological and muscular syndromes caused by dominant mutations in Ant1 (37–39).

## Results

### Super-toxic Aac2 mutants dominantly kill cells

We previously showed that four pathogenic Ant1 variants modeled in the yeast Aac2 (Fig. 1A) share numerous dominant phenotypes including cold sensitivity, mitochondrial DNA (mtDNA) instability, a propensity to misfold inside mitochondria, and hypersensitivity to low Δψ conditions (40–42). We reasoned that if the mutant proteins share a common mechanism of toxicity that drives these phenotypes, then combining mutations into a single protein may enhance toxicity. To test this, we transformed the wild-type M2915-6A yeast strain with centromeric plasmids expressing wild-type, single and double mutant *aac2* alleles, and selected for Ura^+^ transformants on glucose medium. We found that transformants expressing *aac2^M114P,A128P^* and *aac2^M114P,A137D^* formed smaller colonies on the selective medium at 25°C relative to wild-type *AAC2* and single mutant alleles (Fig. 1B). Strikingly, transformants expressing *aac2^A106D,M114P^*, *aac2^A106D,A128P^*, *aac2^A106D,A137D^* and *aac2^A128P,A137D^* were unable to form visible colonies at 25°C. Growth of transformants expressing some of the double mutants were improved at 30°C (Fig. S1A). These data suggest that combining missense mutations into a single Aac2 protein increases toxicity, even when expressed from a centromeric vector.

**Fig. 1.**
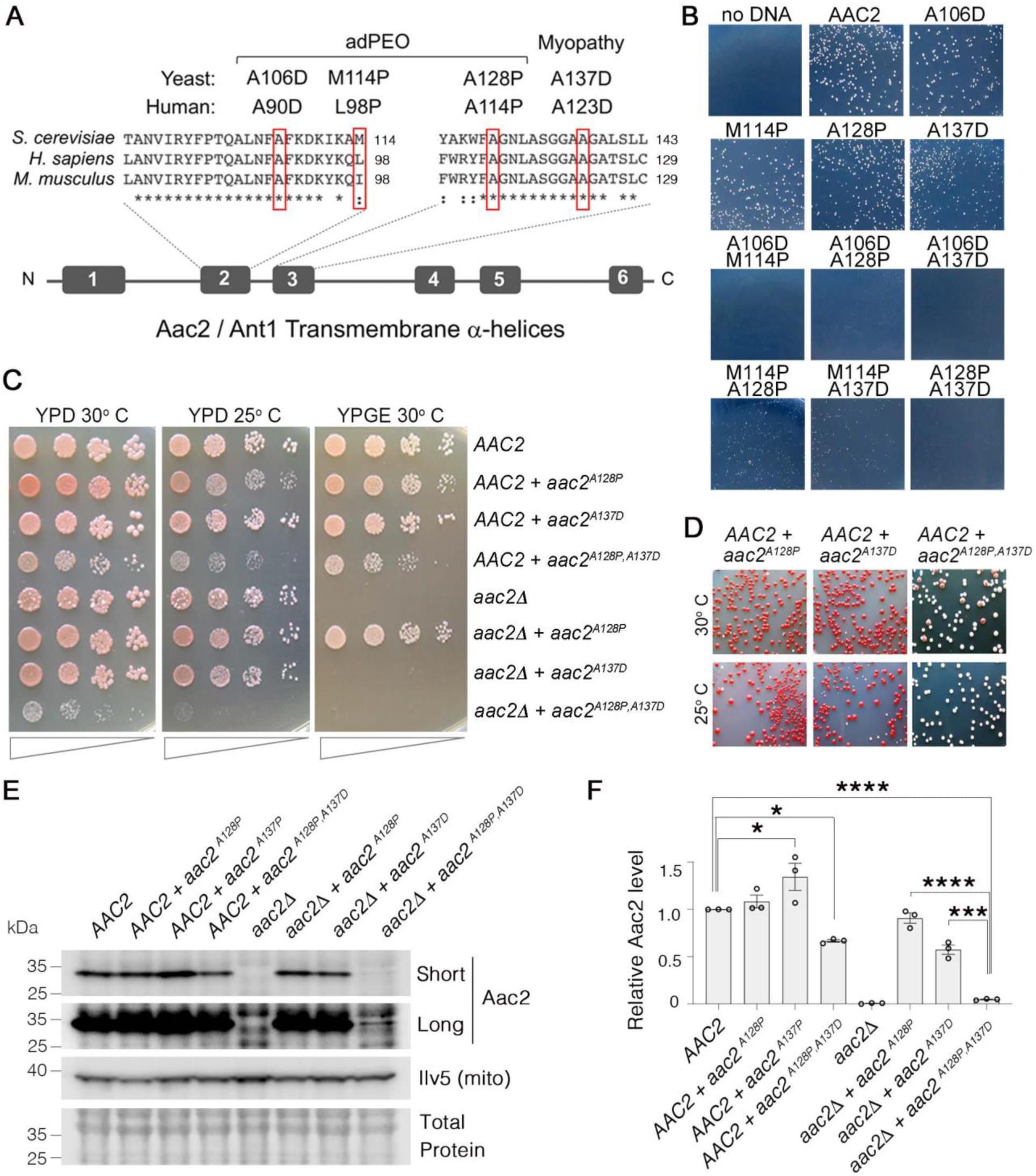
Super-toxic Aac2 mutants dominantly kill cells. **(A)** Schematic showing the location of pathogenic mutations in transmembrane α-helices 2 and 3 of human Ant1 compared with mouse Ant1 and yeast Aac2. adPEO, autosomal dominant Progressive External Ophthalmoplegia. **(B)** Expression of double mutant *aac2* alleles is highly toxic. The yeast M2915-6A strain was transformed with the centromeric vector pRS416 (*URA3*) expressing wild-type or mutant *aac2* alleles and transformants were grown on selective glucose medium lacking uracil at 25°C for 3 days. **(C)** Growth of yeast cells after serial dilution, showing dominant toxicity of *aac2^A128P,A137D^* that is integrated into the genome in the W303-1B strain background. YPD, yeast peptone dextrose medium; YPGE, yeast peptone glycerol ethanol medium. **(D)** The *aac2^A128P,A137D^* allele dominantly increases the frequency of “petite” colonies, which are white. This indicates mtDNA destabilization. **(E)** Immunoblot analysis showing extremely low levels of Aac2^A128P,A137D^. Ilv5 was used as a loading control for mitochondrial protein. Total protein determined with Total Protein Stain (LI-COR). Short, short exposure; Long, long exposure. **(F)** Quantitation from three independent experiments shown in (E). Aac2 values were normalized by Ilv5 to control for mitochondrial content, and data were represented as relative to wild-type; * indicates *p* <0.05, *** *p* <0.001, **** *p* < 0.0001 from one-way ANOVA with Tukey’s multiple comparisons test. Data represented as mean +/- SEM.

We integrated a single copy of *aac2^A128P,A137D^* into the genome of both *AAC2* and *aac2Δ* strains in the W303-1B background. We chose this strain background for two reasons: first, it is more tolerant of mutant *aac2* expression (40); and second, mutant *aac2* expression is not ρ^0^-lethal in this background (like it is in M2915-6A and BY4741), meaning cells expressing mutant *aac2* alleles can survive the loss of mtDNA. This allows us to score mtDNA instability via quantitation of smaller white colonies (“petites”) that form when mtDNA is depleted (43). We found that growth of cells co-expressing *aac2^A128P,A137D^* and *AAC2* is reduced on glucose medium (Fig. 1C), and they form petites at a much higher frequency compared with those expressing the *aac2^A128P^*and *aac2^A137D^* single mutant alleles (Fig. 1D). In the *aac2Δ* background, neither Aac2^A137D^ nor Aac2^A128P,A137D^ supported respiratory growth, consistent with the A123D/A137D mutation eliminating nucleotide transport activity (44). Cells expressing only *aac2^A128P,A137D^* were barely viable, and cell growth was completely inhibited at 25°C even on glucose medium (Fig. 1C). This is in sharp contrast to the *AAC2*-null strain.

Interestingly, we found that Aac2^A128P,A137D^ accumulates to just 4.7% of wild-type Aac2 levels (Fig. 1E-F). We did not observe any accumulation of aggregated Aac2^A128P,A137D^ (Fig. S1B-C), indicating that Aac2^A128P,A137D^ degradation (see below), rather than aggregation, is the likely explanation for low protein recovery. Taken together, the data indicate that *aac2^A128P,A137D^* imparts severe toxicity through a mechanism that is independent of nucleotide transport.

### Super-toxic Aac2 proteins clog the TOM complex

We hypothesized that the highly toxic double mutant Aac2 is arrested during mitochondrial protein import thereby clogging protein import and impairing mitochondrial biogenesis. To test this, we imported ^35^S-labeled Aac2 variants into wild-type mitochondria and analyzed the import reaction by blue native polyacrylamide gel electrophoresis (BN-PAGE) followed by autoradiography (Fig. 2A) (18, 45). We did not observe a significant reduction in the amount of IMM-inserted “mature” Aac2^A128P^ or Aac2^A137D^ compared with wild-type Aac2, suggesting no obvious import defect when wild-type mitochondria are fully energized *in vitro*. In contrast, integration of the double mutant Aac2^A128P,A137D^ into the IMM was reduced by >70% (Fig. 2A-B). To determine whether the preprotein can enter mitochondria, we treated the import reaction with proteinase K to digest non-imported preproteins. We found that a significant portion of mutant Aac2, particularly Aac2^A128P,A137D^, is sensitive to proteinase K digestion (Fig. 2C-D). These data suggest that either the Aac2^A128P,A137D^ preprotein is not transported to the TOM complex, or it fails to traverse the Tom complex. To decipher between these possibilities, we imported Aac2 variants into mitochondria containing hemagglutinin (HA)-tagged Tom40 for affinity purification of the TOM complex. Indeed, Aac2^A128P,A137D^ had increased association with Tom40-HA compared with wild-type (Fig. 2E-F). These observations suggest that a significant fraction of Aac2^A128P,A137D^ is arrested at the TOM complex during import into fully energized wild-type mitochondria.

**Fig. 2.**
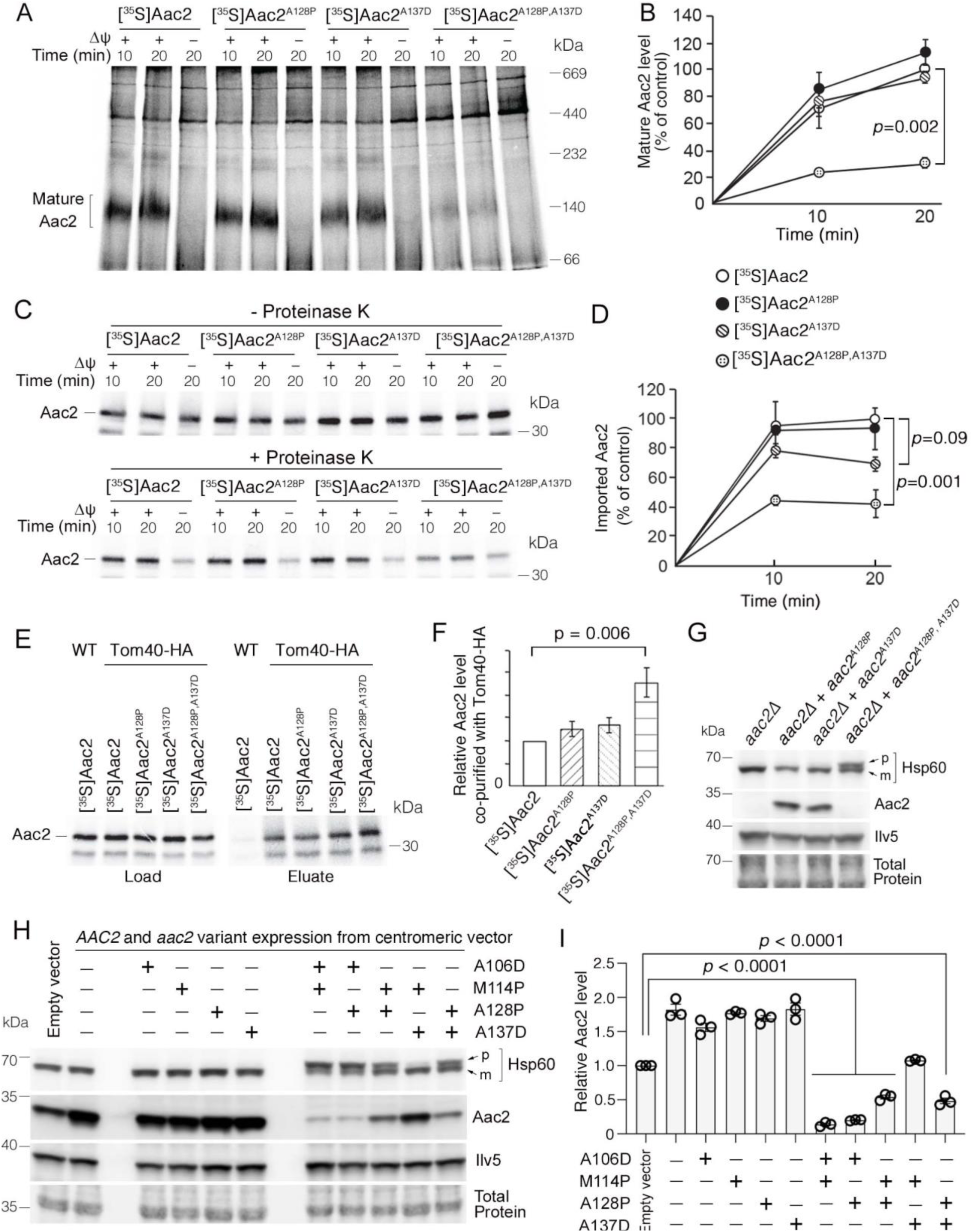
Super-toxic Aac2 proteins clog the TOM complex. **(A)** *In vitro* protein import assay. ^35^S-labeled Aac2 and mutant variants were imported into wild-type mitochondria for 10 or 20 minutes and analyzed by blue native electrophoresis and autoradiography. **(B)** Quantitation from three independent experiments depicted in (A). *P* value from two-way repeated measures ANOVA with Sidak’s multiple comparisons test. **(C)** ^35^S-labeled Aac2 and mutant variants were imported into wild-type mitochondria without (upper) or with (lower panel) subsequent proteinase K treatment to degrade non-imported preproteins. Reaction analyzed by SDS-PAGE and autoradiography. **(D)** Quantitation from three independent experiments depicted in (C). *P* values were calculated as in (B). **(E)** Preferential association of mutant Aac2 with Tom40-HA. ^35^S-labeled Aac2 and mutant variants were imported into Tom40-HA mitochondria, followed by anti-HA immunoprecipitation and analysis by SDS-PAGE and autoradiography. **(F)** Quantitation from three independent experiments depicted in (E). *P* value was calculated with a one-way ANOVA and Dunnett’s multiple comparisons test. **(G)** Immunoblot analysis showing accumulation of the un-cleaved Hsp60 precursor (p) in cells expressing chromosomally integrated aac2^A128P,A137D^. Cells were grown in YPD at 30°C. m, mature (i.e. cleaved). **(H)** Immunoblot analysis showing accumulation of un-cleaved Hsp60 precursor (p) in cells expressing aac2 variants from a monocopy/centromeric vector. Cells were grown in YPD at 25°C. **(I)** Quantitation from three replicates of (H). Aac2 values normalized by the mitochondrial protein Ilv5, and then normalized to vector-transformed samples. P values were calculated as in (F). Data represented as mean +/- SEM.

Next we wondered whether Aac2^A128P,A137D^ clogs the TOM complex *in vivo*. If it does, we would expect the accumulation of unprocessed mitochondrial preproteins that travel through the TOM complex, which would include the vast majority of intramitochondrial proteins including both TIM22 and TIM23 substrates. Indeed, precursor of the mitochondrial matrix protein Hsp60 was readily detectable in cells expressing Aac2^A128P,A137D^ despite extremely low levels of the mutant protein (Fig. 2G). We also acutely expressed *aac2* from galactose-inducible *GAL10* promoter to limit possible indirect effects on protein import efficiency. As expected, expression of mutant *aac2* from the G*AL10*-promoter was highly toxic (Fig. S2A). Importantly, the timing of Hsp60 precursor accumulation (Fig. S2B) was coupled with the induction of the mutant Aac2 (Fig. S2C). The data suggest that the mutant Aac2 directly affects protein import *in vivo*.

We extended our analysis to include additional single and double mutant *aac2* alleles (see Fig. 1A). In transformants expressing double mutant *aac2* alleles from a monocopy vector, the level of un-cleaved Hsp60 correlates with toxicity (Fig. 2H, see also 1B and S1A). Moreover, total Aac2 levels were reduced by all double mutant Aac2 variants except the least toxic Aac2^M114P,A137D^ (Fig. 2H-I) suggesting impaired biogenesis of endogenous wild-type Aac2. The extent of endogenous Aac2 reduction also correlated with cell toxicity. These data further support the idea that protein import clogging underlies the super-toxicity of Aac2 double mutant.

### Yme1, but not the proteasome, contributes substantially to Aac2^A128P,A137D^ degradation

We hypothesized that the proteasome should degrade Aac2^A128P,A137D^ as it clogs at the TOM complex. To test this, we first generated strains lacking the drug efflux pump Pdr5, which allows accumulation of the proteasome inhibitor MG132 in yeast cells. Proteasome inhibition by MG132 was confirmed by the accumulation of Sml1, a labile proteasome substrate (Fig. 3A) (46). The experiment was performed in *aac2Δ* background to facilitate Aac2^A128P,A137D^ detection. Interestingly, we found that MG132 failed to increase the steady-state level of Aac2^A128P,A137D^ (Fig. 3A). Secondly, we found that proteasome dysfunction induced by two temperature-sensitive mutants, *ump1Δ* and *pre9Δ* (Fig. 3B), also failed to increase Aac2^A128P,A137D^ levels (Fig. 3C). Finally, cycloheximide chase following a galactose-induced Aac2^A128P,A137D^ synthesis confirmed that the mutant protein is unstable compared with the wild-type Aac2. More importantly, MG132 failed to stabilize Aac2^A128P,A137D^ (Fig. 3D-E). These data suggest that little, if any, Aac2^A128P,A137D^ is degraded by the proteasome.

**Fig. 3.**
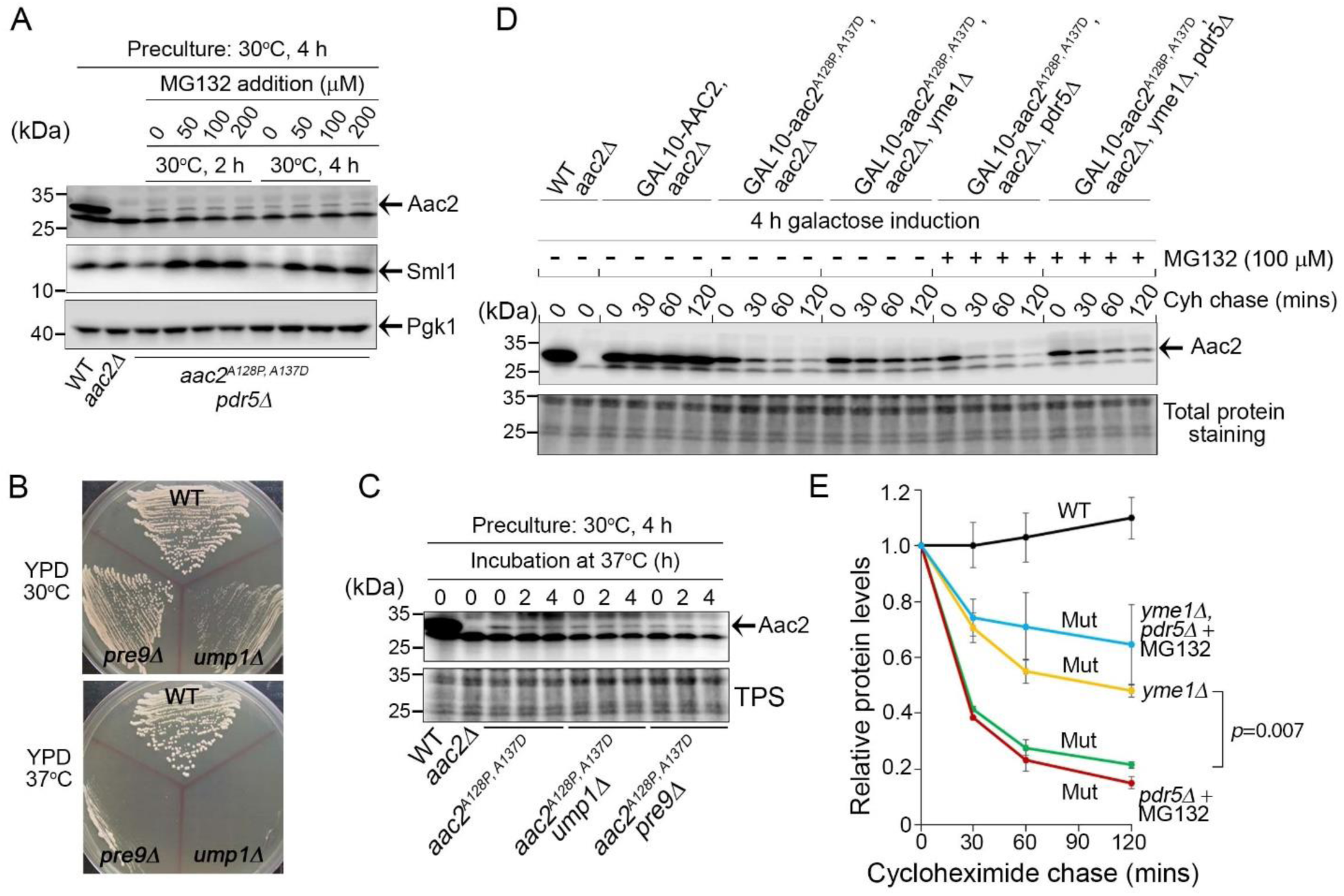
Degradation of Aac2^A128P, A137D^ by Yme1. **(A)** Effect of MG132 on the steady state level of Aac2^A128P, A137D^ in a strain disrupted of *PDR5* and *AAC2*. Cells were first grown in YPD medium at 30°C for four hours before MG132 was added at the indicated concentrations. Cells were cultured for another two or four hours before being harvested for western blot analysis. Sml1 was used as a control for proteasome inhibition. **(B)** Temperature sensitive phenotype of *ump1Δ* and *pre9Δ* cells. Cells were grown at the indicated temperatures for 2 days before being photographed. **(C)** Western blot analysis showing that *ump1Δ* and *pre9Δ* do not affect the steady state level of Aac2^A128P, A137D^. Cells were grown in YPD medium for two and four hours at the restrictive temperature (37°C) before being analyzed for Aac2^A128P, A137D^ levels. TPS, total protein staining. **(D)** Western-blot showing the stability of Aac2^A128P, A137D^ after cycloheximide (Cyh) chase in cells disrupted of *YME1* or treated with MG132, following *GAL10*-induced synthesis of Aac2^A128P, A137D^ in galactose medium at 30°C for 4 hours. **(E)** Quantification of data for the turnover rate of Aac2^A128P, A137D^ (Mut) and its wild-type control (WT) depicted in (D). Aac2 levels were first normalized by total protein stain and then plotted as values relative to time zero. Depicted are mean values +/- S.E.M. from three independent experiments. *P* values were calculated with a two-way repeated measures ANOVA with Tukey’s multiple comparisons test to compare genotypes at time=120 mins.

Next we tested whether an intramitochondrial proteolytic system is responsible for Aac2^A128P,A137D^ degradation. One likely candidate is Yme1, which is an IMS-facing AAA protease whose genetic deletion is synthetically lethal with mutant *aac2* variants (40). Indeed, we found that Aac2^A128P,A137D^ is significantly but not fully stabilized in cells disrupted of *YME1* (Fig. 3D-E). Inhibition of proteasomal function with MG132 does not significantly stabilize Aac2^A128P,A137D^ in *yme1Δ* cells. These data strongly suggest that Yme1 plays an important role in the degradation of Aac2^A128P,A137D^, possibly serving as a quality control mechanism for degrading stalled import substrates in the vicinity of TIM22 and/or on the OMM (47).

### Aac2^A128P^ accumulates along the carrier import pathway and induces protein import stress

The single mutant Aac2 variants did not show significantly reduced import *in vitro* and only a mild accumulation at the TOM complex (Fig. 2A-B, E-F). However, acute overexpression of the single mutant Aac2^A128P^ led to accumulation of the precursor form of HSP60, consistent with clogging of the translocation apparatus (Fig. S2B). To explore an arrest of Aac2^A128P^ at protein translocases, we performed affinity purification Aac2-HIS_6_ and Aac2^A128P^-HIS_6_ followed by a quantitative proteomic comparison of the co-purified proteins (Fig. 4A-B; S3A-C). Numerous proteins preferentially co-purify with Aac2^A128P^-HIS_6_ over Aac2-HIS_6_ (Data S1), which were enriched for chaperones involved in targeting of mitochondrial preproteins through the cytosol (Fig. 4C; S3D-F). Aac2^A128P^ also has increased association with the TOM complex, as well as the translocase of the inner membrane (TIM22) complex that is responsible for the Δψ-dependent insertion of carrier proteins into the IMM (Fig. 4C). The association with Tim22 was confirmed by immunoblotting (Fig. S3G-H). Expectedly, there is no increased association of Aac2^A128P^ with Tim23, a known binding partner of wild-type Aac2 that is not involved in its import (Fig. S3I) (48). Repeat affinity purification, except with double the ionic strength in the binding buffer, reproduced these data (Fig. S4A-E).

**Fig. 4.**
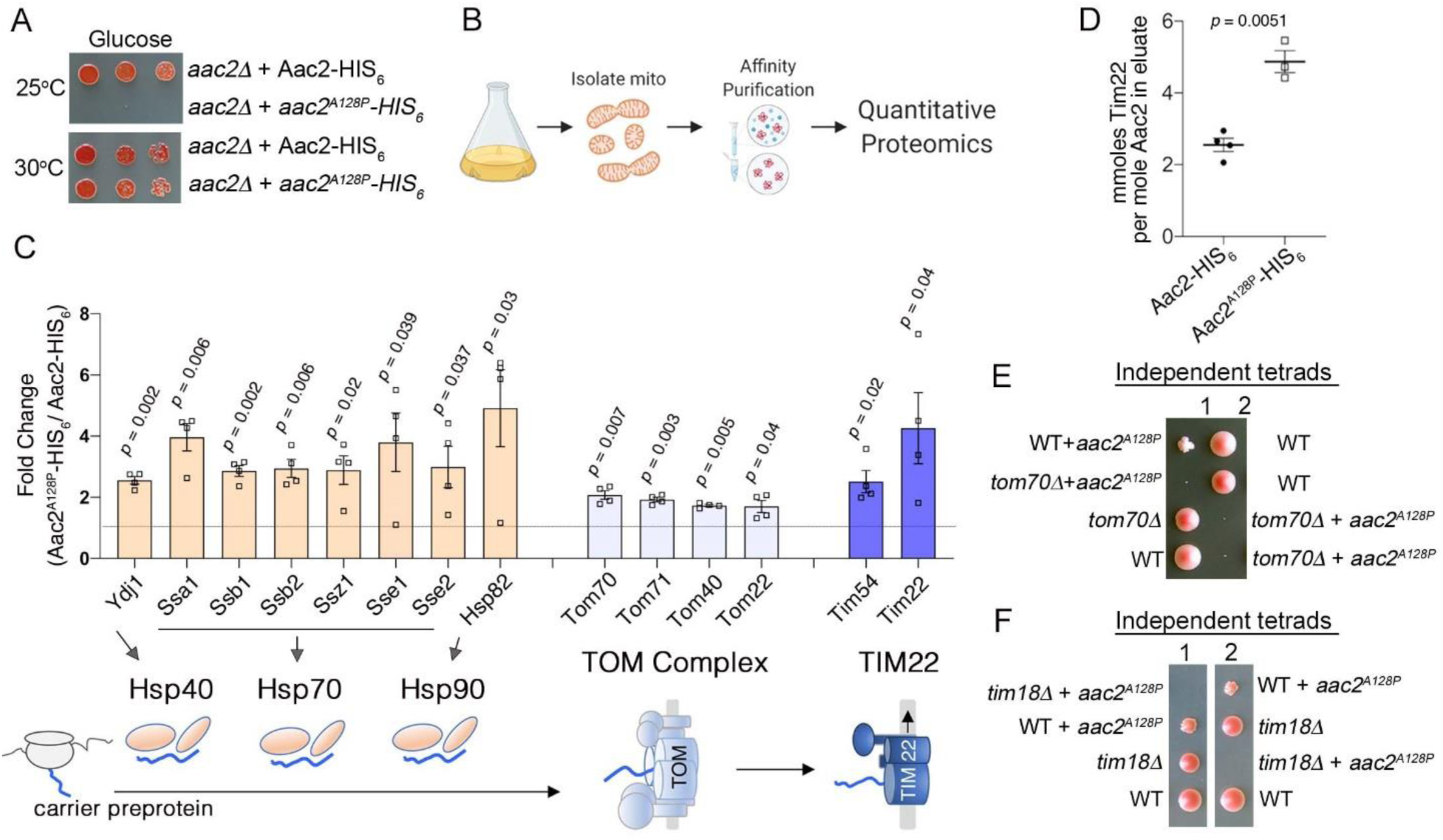
Aac2^A128P^ accumulates along the carrier import pathway and induces protein import stress. **(A)** Growth of cells after serial dilution showing that *aac2^A128P^*-HIS_6_ is toxic at 25°C on glucose medium in a M2915-6A derived strain. **(B)** Schematic of our approach to identify aberrant protein-protein interactions of Aac2^A128P^-HIS_6_. **(C)** Co-purified proteins significantly enriched in Aac2^A128P^-HIS_6_ eluate compared with Aac2-HIS_6_. See Methods for details on abundance value calculation. FDR-corrected *p*-values depicted are from multiple *t* test analysis. Lower panel is a schematic of the mitochondrial carrier protein import pathway. **(D)** Absolute quantities of Aac2 and Tim22 in Aac2-HIS_6_ and Aac2^A128P^-HIS_6_ eluates, as determined by Parallel Reaction Monitoring (PRM) mass spectrometry. *P* value was calculated with student’s *t* test. **(E – F)** Tetrad analysis demonstrating synthetic lethality of *aac2^A128P^* expression with genetic defects in carrier protein import in the M2915-6A strain background. Cell were grown on YPD at 30°C. Data depicted as mean +/- SEM.

To quantitate the increased association of Aac2^A128P^ to Tim22 relative to the wild-type Aac2, we performed targeted quantitative proteomics of Aac2-HIS_6_ pull-down products (Fig. S4F). We found that ∼5 mmoles of Tim22 are pulled down per mole of Aac2^A128P^-HIS_6_ (Fig. 4D). If 5 mmoles per mole of Aac2^A128P^ molecules are bound to Tim22 *in vivo*, this could theoretically occupy ∼85% of Tim22 channels, as Aac2 is present at ∼188,000 molecules per cell and Tim22 at just 1,100 (3). Taken together, the data suggest that Aac2^A128P^ is stalled at both the TOM and TIM22 complexes during import.

Genetic and biochemical analyses provided additional support for a clogging activity associated with Aac2^A128P^. First, we observed that *aac2^A128P^*-expressing cells are sensitive to genetic perturbation to the carrier import pathway. *TOM70* and *TIM18* are two non-essential genes directly involved in the import of mitochondrial carrier proteins such as Aac2, serving as an import receptor at the TOM complex and a component of the TIM22 complex, respectively (10, 26). We found that meiotic segregants combining *aac2^A128P^* expression with *tom70Δ* or *tim18Δ* form barely-visible microcolonies at 30°C on glucose medium, which strongly indicates synthetic lethality (Fig. 4E-F).

Next, we wondered whether the cellular response to acute Aac2^A128P^ expression could be indicative of protein import stress. To test this, we surveyed expression levels of a panel of genes known to be upregulated by clogging the TOM complex. These experiments were performed in the BY4742 strain background, in which expression of *aac2^A128P^* is incompatible with mtDNA loss (Fig. S5A), thus precluding any transcriptional effects of mtDNA destabilization. We found that *CIS1* transcript levels are acutely increased upon expression of *aac2^A128P^*, but not of *AAC2*, from a *GAL10* promoter (Fig. S5B). *CIS1* upregulation is the hallmark of the mitochondrial compromised protein import response (mitoCPR) (49). We also found that *RPN4*, *HSP82*, *SSA3*, and *SSA4* are transiently upregulated (Fig. S5C-F), consistent with previously published gene activation patterns induced by a synthetic mitochondrial protein import clogger (50). Thus, transcriptional responses further suggest that Aac2^A128P^ clogs the protein import machinery.

### Ant1^A114P^ and Ant1^A114P,A123D^ clog mitochondrial protein import in human cells

We introduced equivalent mutations in human Ant1 with a C-terminal hemagglutinin (HA)-tag, and transiently expressed the mutant proteins in HeLa cells (see Fig. 1A). Like in yeast, combining missense mutations in Ant1 dramatically reduced steady-state protein levels, suggesting impaired Ant1 biogenesis (Fig. 5A). Most strikingly, the level of Ant1^A114P,A123D^, equivalent to the yeast Aac2^A128P,A137D^, is only 2.2% of the wild-type. Three lines of evidence suggest that Ant1^A114P^ and Ant1^A114P,A123D^ obstruct protein import into mitochondria. First, immunoprecipitation of Ant1^A114P^-HA demonstrated that the mutant protein has increased interactions with components of both TOM and TIM22 complexes (Fig. 5B-C), as observed with its yeast ortholog Aac2^A128P^. Second, protease protection assay demonstrated that 42% and 52% of total Ant1^A114P^ and Ant1^A114P,A123D^ proteins are exposed on the OMM, whereas the wild-type Ant1 is protected from proteinase K digestion (Fig. 5D-E). This is consistent with *in vitro* import studies of the Aac2 variants in yeast mitochondria (Fig. S2A-D). Third, proteomics demonstrated that mitochondrial proteins accumulate in the cytosol of *Ant1^A114P^* and *Ant1^A114P,^*^A123D^ -transfected cells, compared with wild-type Ant1-transfected cells (Fig. 5F). Moreover, mitochondrial matrix proteins were significantly enriched in the cytosol of Ant1^A114P,A123D^-expressing cells (FDR < 3 x 10^-6^) (Data S2; Fig. 5G), corroborating that the double mutant impairs protein import. We therefore conclude that Ant1^A114P^ and Ant1^A114P,A123D^ clog protein import, with Ant1^A114P,A123D^ having enhanced clogging activity despite low protein levels.

**Fig. 5.**
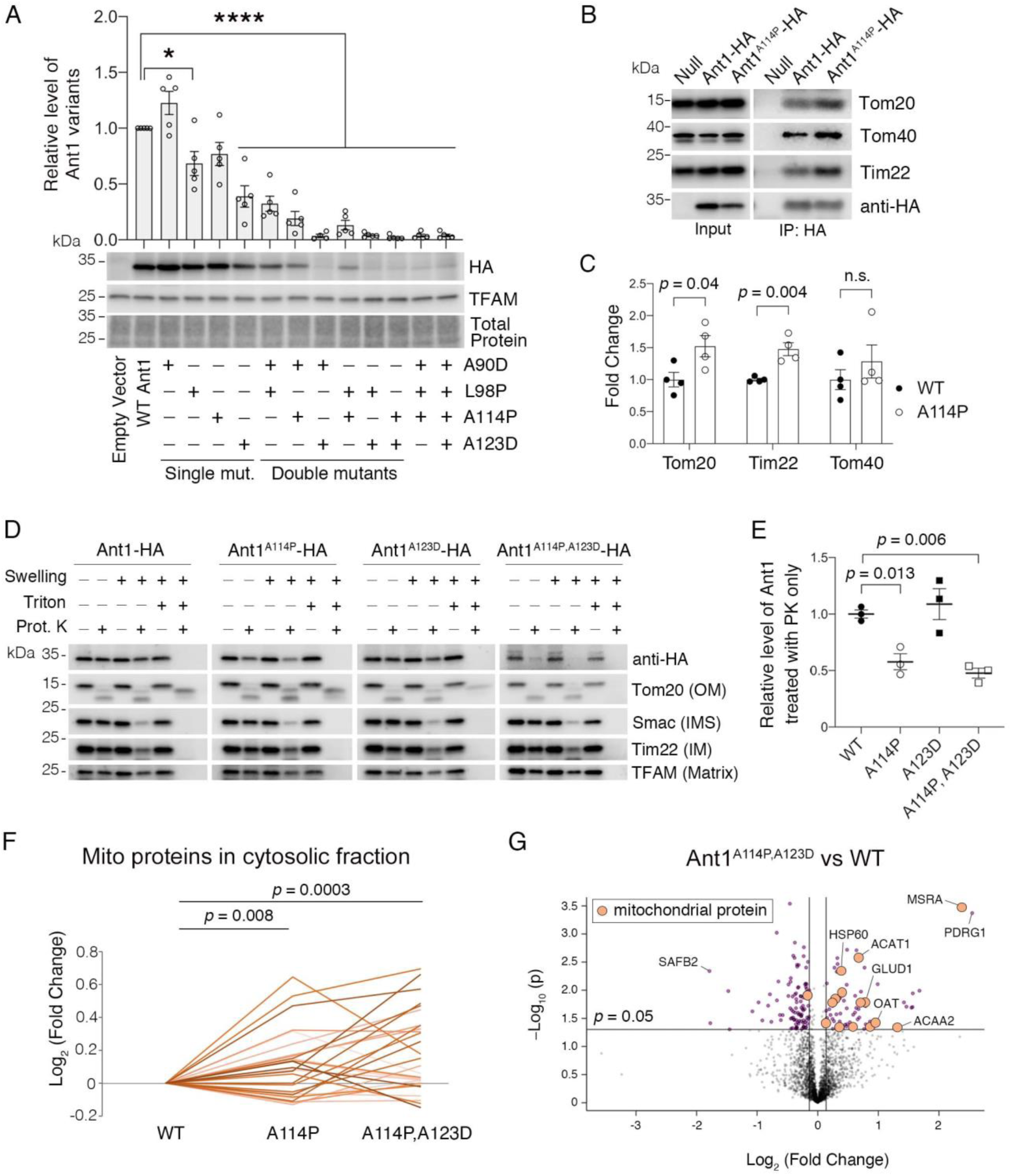
Ant1^A114P^ and Ant1^A114P,A123D^ clog mitochondrial protein import in human cells. **(A)** Combining pathogenic mutations in Ant1 strongly reduces protein levels, as indicated by immunoblot analysis of Ant1-HA levels 24-hours after transfecting HeLa cells. Ant1 variant levels were normalized by TFAM, then plotted as relative to wild-type level. * indicates *p* <0.05, **** *p* < 0.0001 from one-way ANOVA with Dunnett’s multiple comparisons test. **(B)** Immunoprecipitation (IP) of Ant1-HA and Ant1^A114P^-HA from transiently transfected HeLa cells followed by immunoblot analysis, showing that Ant1^A114P^ has increased interaction with the protein import machinery like its yeast ortholog Aac2^A128P^. **(C)** Quantitation from four independent immunoprecipitations, one of which is depicted in (B). *P* values were calculated with a student’s *t* test. **(D)** Immunoblot analysis following protease protection assay showing that Ant1^A114P^ and Ant1^A114P,A123D^ are sensitive to proteinase K (PK) in isolated mitochondria. Swelling in hypotonic buffer was used to burst the outer membrane, and Triton X-100 was used to disrupt all membranes. OM, outer membrane; IMS, intermembrane space; IM, inner membrane. **(E)** Quantitation of the wild-type and mutant Ant1 pools that are protected from PK degradation in intact mitochondria. All HA levels were normalized by TFAM, then plotted as relative to its untreated sample. Replicates from three independent transfections. *P* values were calculated with a one-way ANOVA and Holm-Sidak’s multiple comparisons test. **(F)** Ant1^A114P^ and Ant1^A114P,A123D^ expression obstruct general mitochondrial protein import. Proteomics of the cytosolic fraction of transfected HeLa cells reveals increase in mitochondrial proteins caused by Ant1^A114P^-HA and Ant1^A114P,A123D^-HA expression relative to Ant1-HA. Each line represents a single mitochondrial protein. *P* values were calculated with a student’s *t* test of the average abundance levels of each mitochondrial protein. **(G)** Volcano plot comparing the cytosolic proteome of Ant1^A114P,A123D^ vs Ant1-transfected HeLa cells. Data represented as mean +/- SEM.

An alternative explanation for some of these observations is general mitochondrial damage and/or apoptosis activation leading to reduced Δψ, which would also cause a reduction in protein import. However, neither Ant1^A114P,A123D^ nor its single mutant counterparts reduced Δψ or significantly increased cell death compared with wild-type (Fig. S6A-D). Thus, the effect on protein import by Ant1^A114P^ and Ant1^A114P,A123D^ is likely due to their physical retention in the import pathway rather than due to reduction of Δψ. It is interesting that Ant1^A114P,A123D^ does not increase apoptosis in the immortalized HeLa cells, despite clearly clogging protein import. This may be related to the transient nature of Ant1 expression or an inherent resistance of the immortalized cells to clogging-induced cell death.

### Ant1^A114P,A123D^ causes dominant muscle and neurological disease

We generated a knock-in *Ant1^A114P,A123D^*/+ mouse line to model protein import clogging *in vivo* (Fig. S7A-B). Dominant *Ant1*-induced diseases primarily affect skeletal muscle, with low-penetrant neurological involvement (37–39, 51–53). Key muscle features include mildly reduced mitochondrial respiratory function, COX-deficient muscle fibers, and muscle weakness. Consistent with clinical phenotypes, we found that the maximal respiratory rate (state 3) of Ant1^A114P,A123D^/+ muscle mitochondria was reduced by ∼20% and ∼30% when utilizing complex I in 9- and 24-month old mice, respectively (Fig. 6A). The respiratory control ratio, which is the single most useful and sensitive general measure of energy coupling efficiency (54), was reduced by ∼31 and ∼42% in young and old mice, respectively. Surprisingly, when complex I is inhibited and complex II substrate (succinate) is present, maximal respiration is increased by ∼23% in the mutant mice at 9 months old (Fig. 6B). This result is important because it confirms that ATP/ADP transport is not a limiting factor for sustaining a high respiratory rate in *_Ant1A114P,A123D_*_/+ mice._

**Fig. 6.**
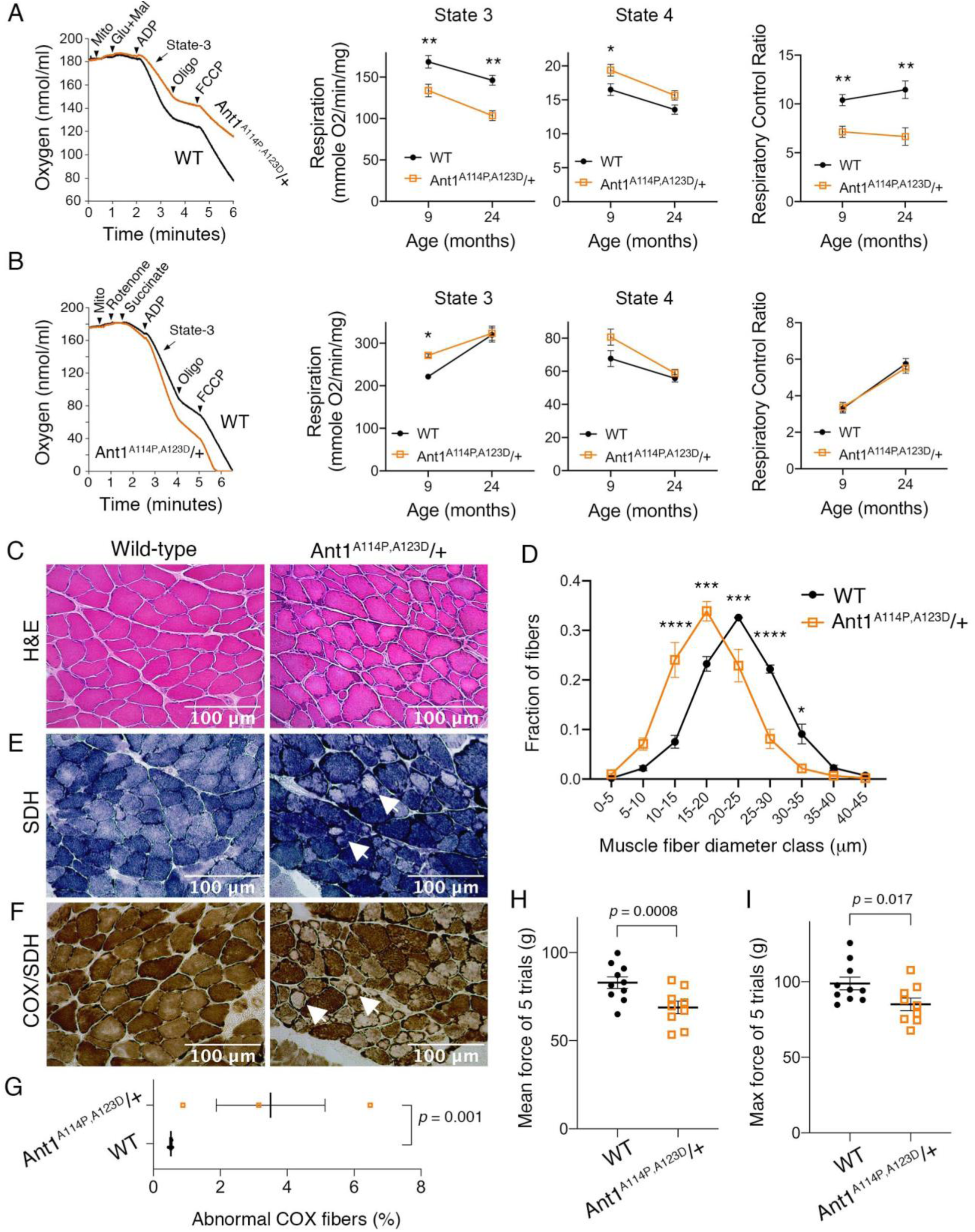
Ant1^A114P,A123D^ causes a dominant mitochondrial myopathy in mice. **(A)** Respirometry of isolated skeletal muscle mitochondria with complex I stimulated by glutamate (glu) and malate (mal). State 3, maximal respiratory rate after addition of ADP; state 4, oligomycin (oligo)-inhibited respiratory rate; Respiratory Control Ratio = State 3 / State 4. N = 6 mice/genotype at 9-months-old; n=4 mice per genotype at 24 months old. 3 measurements were taken per mouse. *P* values were derived from repeated measures ANOVA with measurement order as the within-subjects variable. Data from two age groups were analyzed independently. FCCP, Trifluoromethoxy carbonylcyanide phenylhydrazone. **(B)** Respirometry of isolated skeletal muscle mitochondria with complex II stimulated by succinate and complex I inhibited by rotenone. N = 2 mice/genotype at 9-months-old, 4 measurements/mouse; n = 4 mice/genotype at 24-months-old, 3 measurements/mouse. Data analyzed as in (A). **(C)** Soleus muscles stained with haematoxylin and eosin (H&E) showing smaller myofibers in 30-month old Ant1^A114P,A123D^/+ mice. **(D)** Feret’s diameter analysis of H&E stained soleus in (C) reveals atrophy in Ant1^A114P,A123D^/+ mice. At least 340 myofibers were measured per soleus. Myofiber diameters were binned into 5 μm ranges and plotted as % of total. N = 3 mice/genotype. Data analyzed by two-way ANOVA with Sidak’s multiple comparisons test. **(E)** Succinate dehydrogenase (SDH) histochemical activity staining of the soleus showing abnormal fibers that stain for SDH peripherally but are pale internally (arrows). **(F)** Histochemical cytochrome *c* oxidase (COX) and SDH sequential staining of the soleus shows abnormal fibers that stain for COX peripherally, but do not stain for COX or SDH internally. **(G)** Quantitation of abnormal COX fibers shown in (F). *P* value was calculated from student’s t test. **(H)** Forelimb grip strength is reduced in 30-month-old Ant1^A114P,A123D^/+ mice. *P* value from student’s *t* test. **(I)** Maximal forelimb grip strength is reduced in 30-month-old Ant1^A114P,A123D^/+ mice. *P* value from student’s *t* test. Data represented as mean +/- SEM.

In addition to mild bioenergetic defect, we detected significant reduction in myofiber diameter in the skeletal muscle from aged *Ant1^A114P,A123D^*/+ mice (Fig. 6C-D). Sequential COX/SDH histochemical assay failed to detect any COX-negative/SDH-positive fibers in the aged *Ant1^A114P,A123D^*/+ muscles. Instead, we found that ∼3.5% of myofibers have reduced COX and SDH activity centrally, but remained COX and SDH-positive in the periphery (Fig. 6E-G). Finally, we found that the *Ant1^A114P,A123D^*/+ mice display muscle weakness (Fig. 6H-I). These data demonstrate that *Ant1^A114P,A123D^* induces a dominant myopathic phenotype.

We also observed a neurodegeneration phenotype that culminates in paralysis in some *Ant1^A114P,A123D^*/+ mice (Fig. S7C, Movies S1 & S2). This phenotype occurred in only four Ant1^A114P,A123D^/+ mice, with a penetrance of 3.4% among the heterozygous mice that have reached 15 months old. The presenting symptom is typically altered gait after the age of 11 months, followed by weight loss and death within 2-3 weeks of symptom onset. Histological analysis of the lumbar spinal cord in a symptomatic *Ant1^A114P,A123D^*/+ mouse demonstrated dissolution of Nissl substance in the cell bodies of ventral horn neurons (Fig. S7D), consistent with motor neuron degeneration (55). Neurodegeneration in the spinal cord was also indicated by GFAP accumulation by immunofluorescence and immunoblotting (Fig. S7E-G). Transmission electron microscopy of ventral horn neurons revealed loss of cristae density of mitochondria, which suggests defects in mitochondrial biogenesis (Fig. S7H).

We found that Ant1^A114P,A123D^ is highly unstable in mouse tissues, consistent with yeast and human cell data. We were unable to detect the mutant protein by Western blot in tissue lysate from homozygous mice (Fig. 7A). Only in isolated skeletal muscle mitochondria were we able to detect Ant1^A114P,A123D^, which was present at ∼0.1% of wild-type level (Fig. 7B-C). Protease protection assay on isolated muscle mitochondria demonstrated that Ant1^A114P,A123D^ is more sensitive to proteinase K digestion compared with wild-type Ant1 (Fig. 7D-E), consistent with import clogging *in vivo*. Moreover, Smac also appears to be more sensitive to proteinase K in Ant1^A114P,A123D^/Ant1^A114P,A123D^ mitochondria, while Mdh2 and Tim23 does not (Fig. S7I-K). While this could indicate instability or partial rupture of the OMM in mutant mitochondria, it could also be a result of clogging and imply that clogging does not affect all mitochondrial preproteins equally. Consistent with this concept, our experiments in yeast showed minimal effects of clogging on Ilv5, a mitochondrial matrix protein, in contrast to Hsp60 (Fig. 1E; 2H-I).

**Fig. 7.**
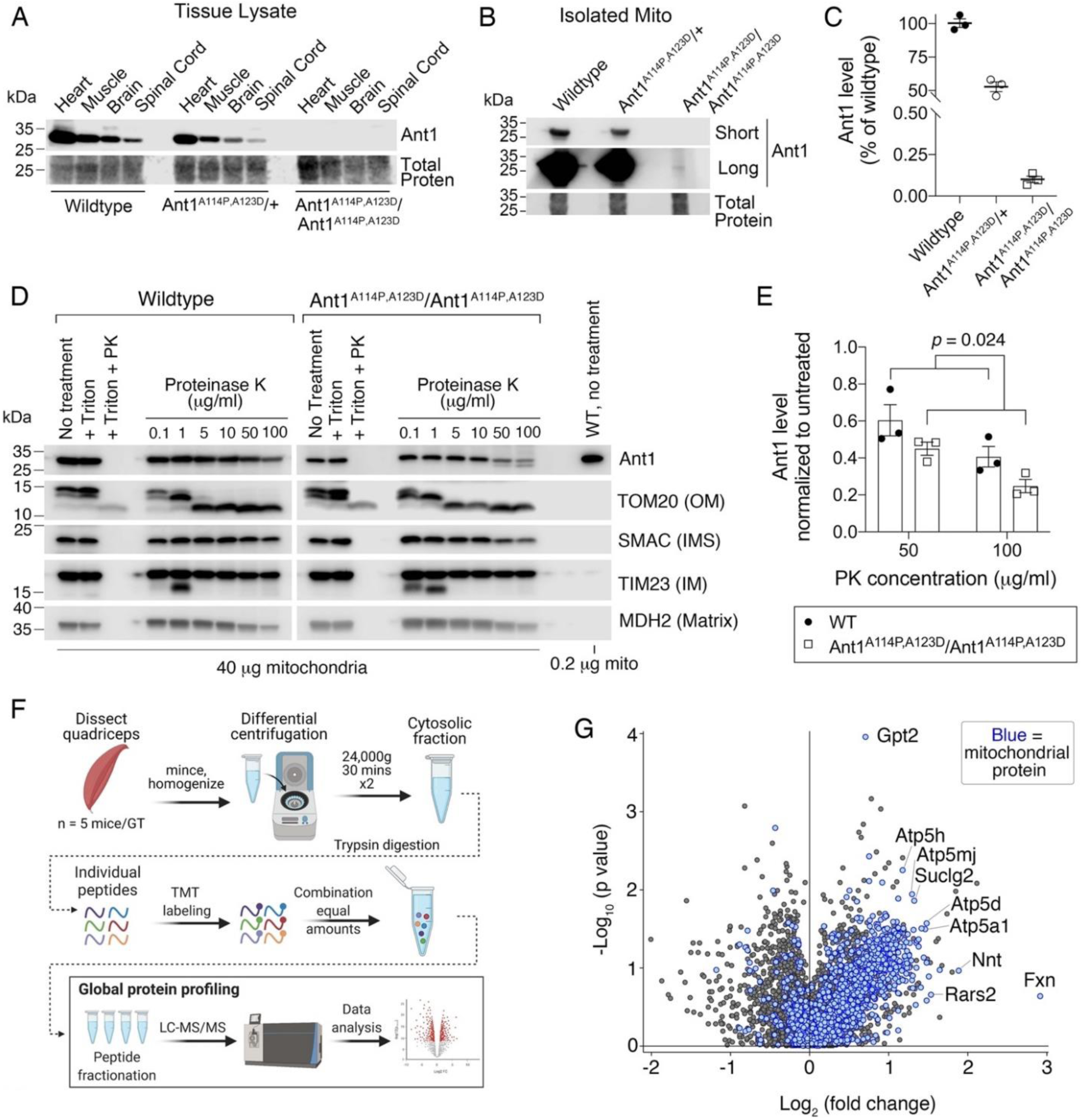
Ant1^A114P,A123D^ clogs protein import *in vivo*. **(A)** Immunoblot analysis of tissue lysate showing low Ant1^A114P,A123D^ protein levels in heterozygous and homozygous mice. **(B)** Immunoblot analysis of isolated muscle mitochondria demonstrating low Ant1^A114P,A123D^ protein levels. **(C)** Quantitation of Ant1 levels in isolated muscle mitochondria from three mice per genotype, as determined by immunoblotting. Values were normalized to total protein stain and shown as relative to wild-type. **(D)** Ant1^A114P,A123D^ is more sensitive to proteinase K (PK) than wild-type Ant1 in intact mitochondria. Immunoblot analysis after PK protection assay of isolated muscle mitochondria in isotonic buffer. Ant1^A114P,A123D^ was detected using SuperSignal West Femto Maximum Sensitivity Substrate (top right panel). **(E)** Quantitation from protease protection assay, as shown in (D). n = 3 mice per genotype. *P* value was calculated with a two-way ANOVA, showing a significant main effect of genotype. Data represented as mean +/- SEM. **(F)** Schematic of Tandem Mass Tagged (TMT) quantitative proteomic analysis. **(G)** Volcano plot comparing the cytosolic proteome of Ant1^A114P,A123D^/+ vs wild-type skeletal muscle, with mitochondrial proteins highlighted in blue.

Finally, to test if Ant1^A114P,A123D^ obstructs general mitochondrial protein import *in vivo*, we compared the cytosolic proteomes of aged muscle from wild-type and *Ant1^A114P,A123D^*/+ mice using Tandem Mass Tagged (TMT) quantitative proteomics (Fig. 7F). We found a striking global increase in mitochondrial proteins in the cytosol of Ant1^A114P,A123D^/+ muscle (Fig. 7G). Among the 75 proteins increased by at least 25% in the cytosol (*p* < 0.05), proteins assigned to the mitochondrion accounted for 45 of them (Data S3). This enrichment was highly significant (FDR < 10^-33^). Taken together, the data suggest that Ant1^A114P,A123D^ clogs general protein import *in vivo*, which may contribute to skeletal muscle pathology by disrupting cytosolic proteostasis.

In summary, we found that Ant1^A114P,A123D^ dominantly causes muscle and low-penetrant neurological disease in mice that recapitulates many pathological and molecular phenotypes of dominant Ant1-induced diseases in humans. This corroborates the toxicity of Aac2^A128P,A137D^ in yeast (Fig. 1), and supports the idea that mitochondrial protein import clogging by a substrate preprotein is pathogenic.

## Discussion

Failure in mitochondrial protein import has severe physiological consequences. In addition to defective mitochondrial biogenesis and energy metabolism, it also causes toxic accumulation and aggregation of mitochondrial preproteins in the cytosol, a process termed mitochondrial Precursor Overaccumulation Stress (mPOS) (56–58). To prevent these consequences, many cellular safeguards have been identified that can maintain protein import efficiency and/or mitigate mPOS (49, 50, 59–73). So far, studies on mitochondrial stressors that impair protein import have been mainly focused on mutations that directly affect the core protein import machinery, or on pharmacological interventions that reduce Δψ (58). In this report, we have uncovered the first example of naturally occurring missense mutations in an endogenous mitochondrial protein causing toxic protein import clogging. We showed that a single protein without overexpression is sufficient to clog import, thereby inducing robust protein import stress responses, bioenergetic defects, and cytosolic proteostatic stress. Ultimately, this import clogging kills yeast cells and causes muscle and neurological degeneration in mice.

### Mitochondrial protein import can be clogged by a mutant mitochondrial preprotein

The mitochondrial carrier protein family is the largest of the transporter families and has highly conserved domain and sequence features across all eukaryotes. Their translocation through the TOM and TIM22 complexes is thought to require partially folded conformations called ‘hairpin loops’ that place adjacent hydrophobic transmembrane α-helices parallel with one another (9, 14). As such, introducing a proline or aspartic acid into the α-helices of Ant1/Aac2 (see Fig. 1A) may prevent hairpin loop formation, rendering the protein incompatible with efficient transit through TOM and TIM22. It is also possible that the proline and aspartic acid affect binding to the Tim9-10 complex, which binds the hydrophobic transmembrane α-helices of preproteins in a manner that seems to include residual α-helical structure (17, 74). The common principle between these possibilities is a loss of the preprotein’s biophysical compatibility with one or multiple binding partners during protein import. This incompatibility not only affects import of the mutant preprotein itself, but also obstructs other preproteins that compete for the same import machinery. Ultimately this reduces cell viability through multi-pronged effects including respiratory defects and cytosolic proteotoxicity.

One of the most surprising observations of this study was the extreme toxicity of the double mutant clogger proteins. The double mutant variants have increased arrest at the TOM complex, which likely reflects severe distortion of their hairpin loops as each mutation occurs in transmembrane α-helices (see Fig. 1A). The deleterious stressors downstream of clogging are likely to be multifactorial and include respiratory defects and cytosolic protestatic stress via mPOS. While it is possible that general proteostasis is perturbed by Aac2^A128P,A137D^ saturating proteolytic machinery, our data suggest this is unlikely. The IMM-associated AAA protease Yme1 (and not the proteasome) appears to be a major proteolytic pathway for clogged Aac2^A128P,A137D^, consistent with the observation that Yme1 may play a role of protein quality control in the vicinity of TIM22 (75, 76). That genetic deletion of *YME1* is well-tolerated in yeast cells suggests that loss of Yme1-based proteolysis does not underlie toxicity of *aac2^A128P,A137D^* expression. The data presented in this study suggest that clogging of the TOM complex is the primary mechanism of Aac2^A128P,A137D^-induced protein import defects and the ensuing cell and organismal toxicity. Importantly, this is strongly supported by *in vitro* experiments showing that Aac2^A128P,A137D^ is defective in translocating through the TOM complex in fully-energized, wild-type mitochondria.

In the context of low Aac2^A128P,A137D^ levels in cells (just 4.7% of wild-type), it is important to note that the adenine nucleotide translocator is one of the most abundant proteins in mitochondria, outnumbering the Tom40 pore-forming subunit of the TOM complex by almost an order of magnitude (3, 77). Thus, if ∼70% of the reduced level of Aac2^A128P,A137D^ is actively clogging TOM complexes, this would still occupy ∼30% of the Tom40 channels, which may have a considerable effect on general protein import. This reflects the central importance of proper TOM complex function to mitochondrial and cell homeostasis, and also provides physiological justification for the existence of multiple pathways dedicated to unclogging the TOM complex (49, 66).

### Mitochondrial protein import clogging as a mechanism of disease

A long-standing mystery in the mitochondrial disease field has been how dominant mutations in the monomeric Ant1 protein cause such a wide spectrum of clinical and molecular phenotypes (37–39, 44, 52, 53, 78). The phenotypes observed in Ant1^A114P,A123D^/+ mice cannot be explained by haploinsufficiency or a dominant negative effect on ADP/ATP exchange. First, Ant1 likely functions as a monomer, thus a dominant-negative mechanism is unlikely (79–81). Second, Ant1^A114P,A123D^’s yeast ortholog, Aac2^A128P,A137D^, is catalytically inactive and clearly exerts toxicity even in conditions where *aac2Δ* cells are healthy (Fig. 1C). Third, mitochondria from Ant1^A114P,A123D^/+ mice are able to exceed the maximal respiration rate of wild-type mitochondria when complex II is stimulated (Fig 6B), indicating that one functional copy of Ant1 is sufficient to support high respiratory rates. Fourth, humans and mice heterozygous for a nonfunctional copy of *Ant1* do not develop neurological or muscle disease (44, 82, 83), while *Ant1^A114P,A123D^*/+ mice develop both. Thus, both the muscle and neurodegenerative phenotypes in *Ant1^A114P,A123D^*/+ mice occur independent of nucleotide transport.

In human and yeast cells, the autosomal dominant Progressive External Ophthalmoplegia (adPEO)-causing Ant1^A114P^ protein (and Aac2^A128P^ in yeast) clearly clogs global mitochondrial protein import, suggesting a new molecular explanation for dominant Ant1-induced disease. Clogging was drastically enhanced by the addition of an A123D (A137D in Aac2) mutation, and the main clogging site shifted to the TOM complex. We note that this mutation was discovered in a homozygous myopathy patient, and that it eliminates nucleotide transport activity (44). Toxicity of Ant1^A114P,A123D^ is therefore independent of nucleotide transport activity. Ant1^A114P,A123D^ expression in mice phenocopies adPEO patients, as Ant1^A114P,A123D^/+ mice exhibit mild OXPHOS defects, partially COX-deficient myofibers, moderate myofiber atrophy, and muscle weakness (51, 84). The low-penetrant neurodegeneration in Ant1^A114P,A123D^/+ mice may also reflect human disease process, as a neurodegenerative phenotype was recently reported in an adPEO patient carrying the *Ant1^A114P^* allele (39). Taken together, these data strongly argue that protein import clogging contributes to adPEO in humans, despite Ant1^A114P,A123D^ not directly genocopying a particular clinical condition.

In summary, we demonstrate that pathogenic mutations in a mitochondrial carrier protein can cause clogging of the protein import pathway, which induces multifactorial cellular stress and disease in mice. These findings uncovered a vulnerability of the mitochondrial protein import machinery to single amino acid substitutions in mitochondrial inner membrane preproteins. There are 53 carrier proteins in human mitochondria that have highly conserved domain organization both within and across species (85). Mutations altering conformational elements involved in protein import, such as hairpin loops, are likely strongly selected against during evolution. Those persisting in the human population would affect protein import and cause disease, as exemplified by the dominant pathogenic mutations in *Ant1*.

Our work provides just a first glimpse into the amino acid requirements of mitochondrial carrier proteins to maintain compatibility with the mitochondrial protein import machinery. Future work will be crucial to systematically evaluate the amino acid requirements first within Ant1/Aac2, and then across entire carrier protein family. It will also be important to determine which specific interactions with protein import components are affected by different mutations. Overall, these efforts may enable the prediction of pathogenic mechanisms of future variants of uncertain significance as they arise with more exome sequencing of human patients.

## Materials and Methods

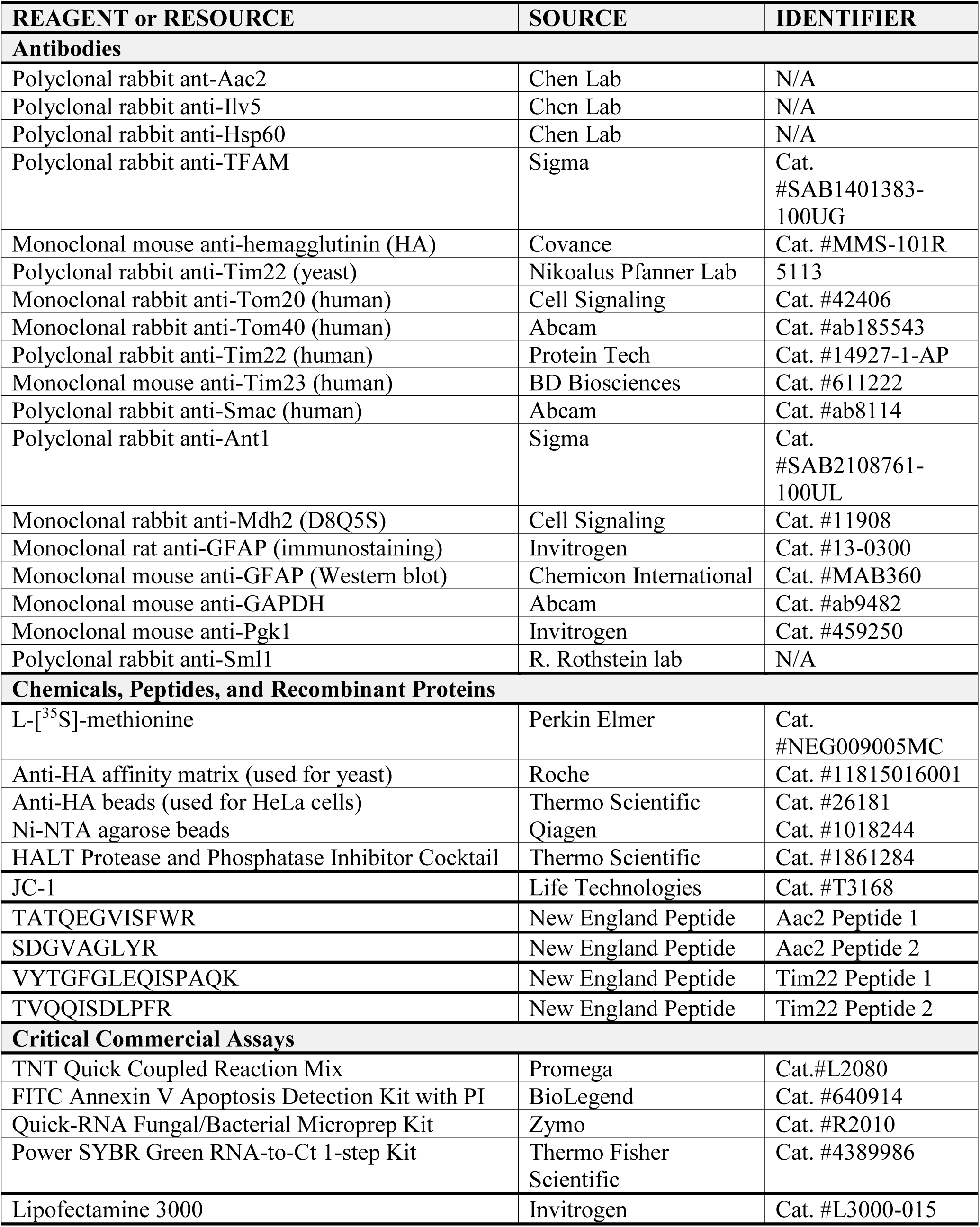
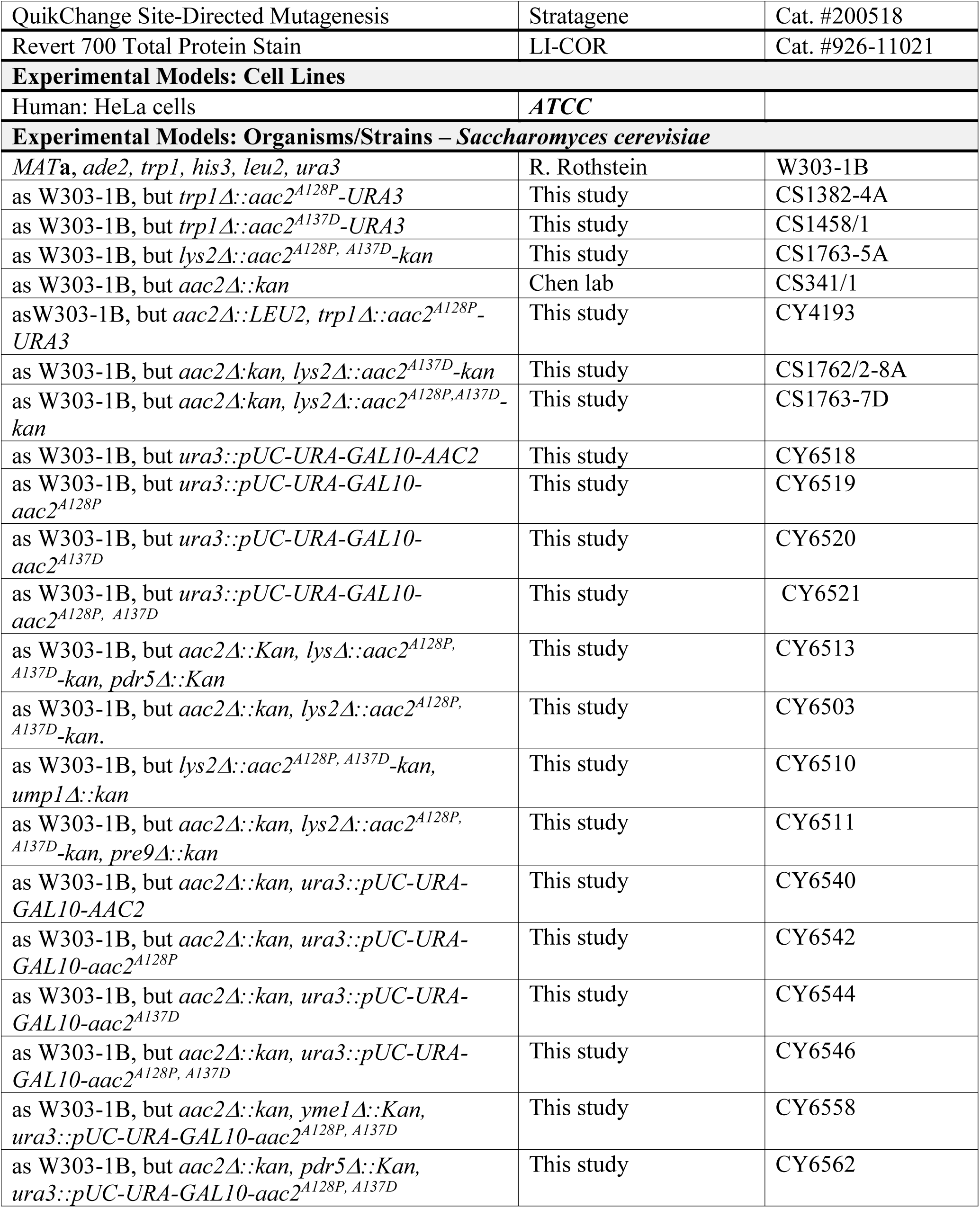
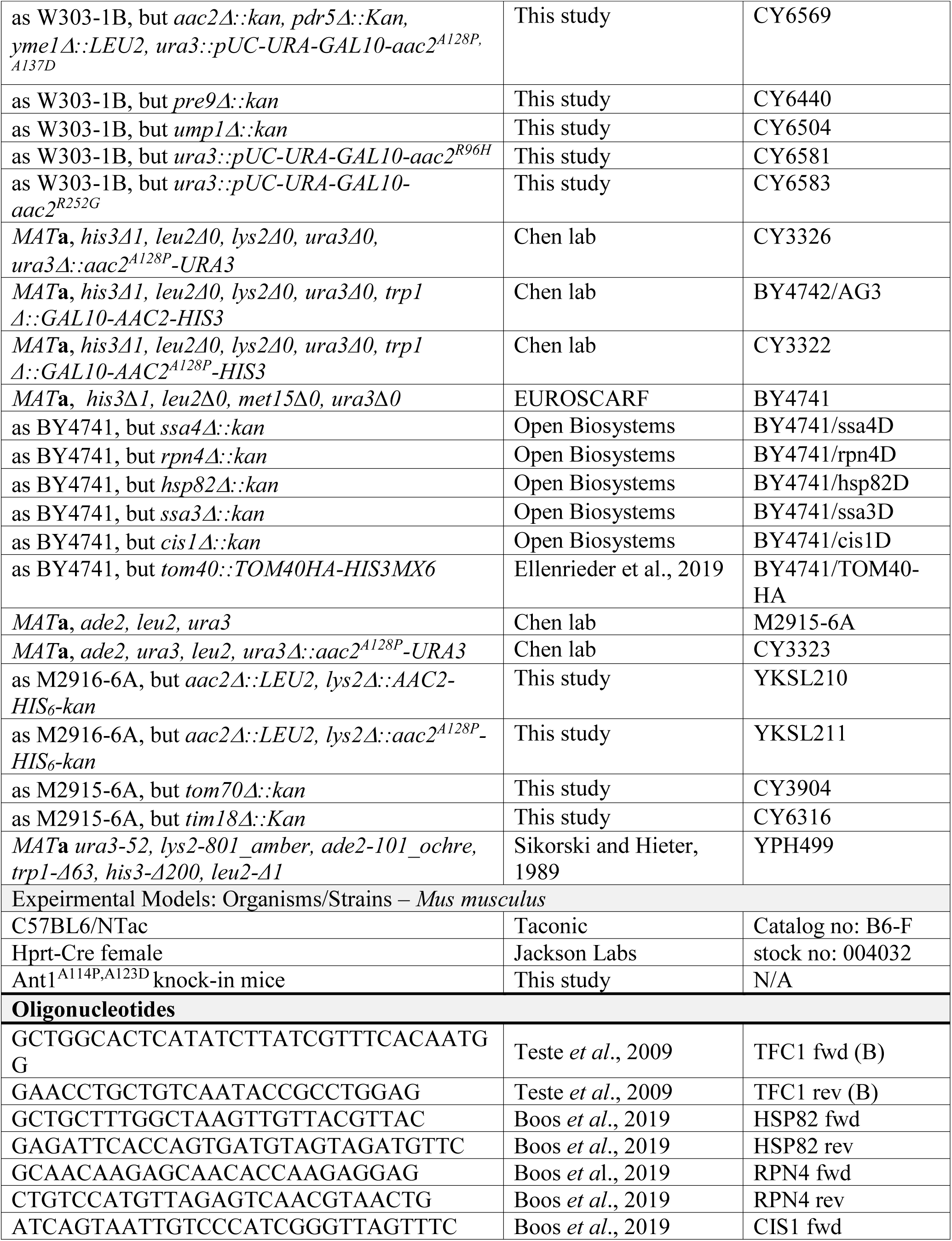
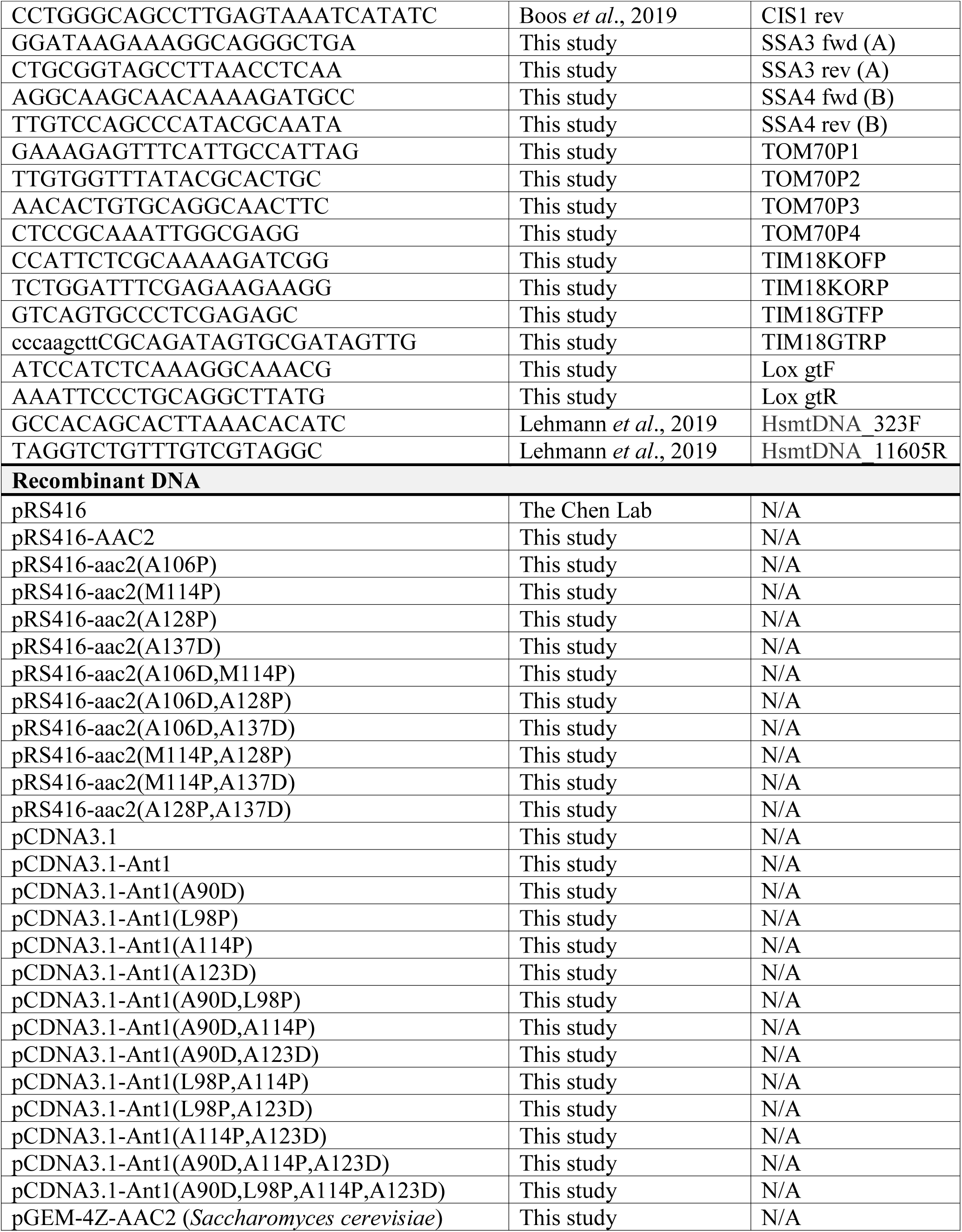
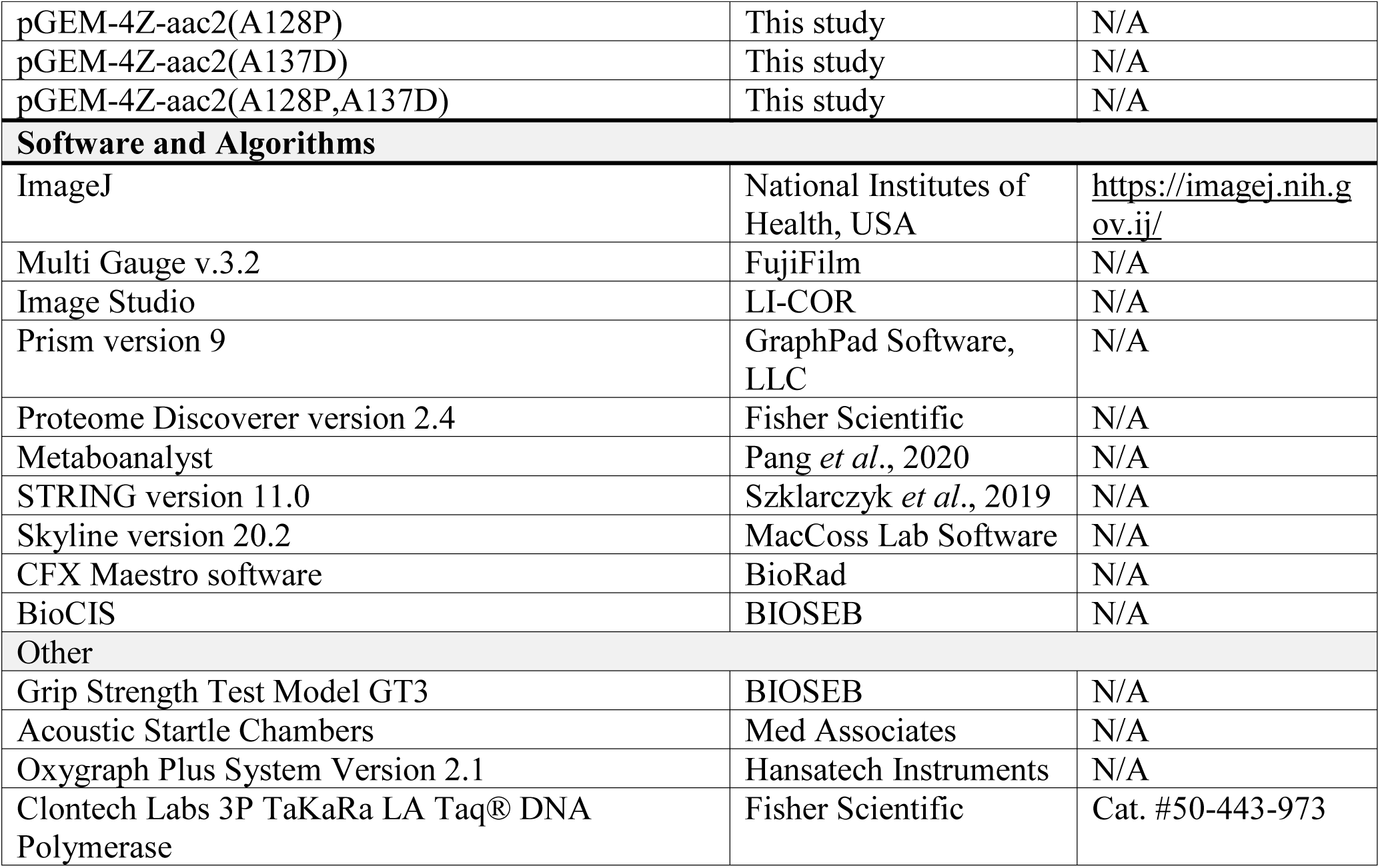

### Yeast growth conditions and genetic manipulation

Yeast cells were grown using standard media. Genotypes and sources of yeast strains are listed in Key Resources Table. To generate the *AAC2* expression plasmids, we amplified the gene from total genomic DNA and inserted it into pFA6a-ScAAC2-URA3-HIS3/2. The missense mutations were introduced using QuikChange Site-Directed Mutagenesis (Stratagene) and mutations were confirmed by sequencing. Mutant genes were then cloned into pRS416 for expression (Fig. 1B) and pFA6a-KanMX6 for placing next to the *KAN* selection marker. The *aac2^A128P,A137^*-*KAN* cassette was then amplified for integration into the *LYS2* locus of the W303-1B strain background. All strains in Fig. 1C-E were derived from this strain by standard genetic crosses. All combinations of *aac2* mutations were expressed by transforming an equal number of M2915-6A yeast with freshly-prepared pRS416-based vectors (*URA3*) expressing wild-type or mutant *aac2*. Equal fractions of the transformant culture were plated on selective medium lacking uracil and grown at 25° or 30°C for 3 days before being photographed (Fig. 1B and S1A). *tom70Δ* and *tim18Δ* strains were generated by amplification of the knockout cassette from genomic DNA of *tom70Δ* and *tim18Δ* strains from BY4741 knockout library, followed by transformation into M2915-6A by selecting for G418^R^. The disruption of the genes was confirmed by PCR using an independent primer pair surrounding the native genomic locus. *PRE9* and *UMP1* were disruption in strains of W303 background by the insertion of *kan*. These alleles were then introduced into strains expressing aac2^A128P, A137D^ by genetic crosses. For Galactose-induced expression, *AAC2* and its mutant alleles were placed under the control of the *GAL10* promoter in the integrative vector pUC-URA3-GAL. The resulting plasmids were then linearized by cutting with *Stu* I within the *URA3* gene, before being integrated into the *ura3*-1 locus by selecting for Ura^+^ transformants. Correct chromosomal integration was confirmed by examining the stability of the Ura^+^ phenotype and PCR amplification of the *URA3* locus. The *yme1Δ* and *pdr5Δ* alleles were introduced into these strains either by direct gene disruption or by genetic crosses.

### Cell lysis, western blotting and signal quantitation

Unless otherwise noted, yeast cells were lysed and prepared for SDS-PAGE as previously described (86). HeLa cells and mouse tissues were lysed with RIPA buffer containing 1X HALT Protease and Phosphatase Inhibitor Cocktail (Thermo Fisher) and prepared for SDS-PAGE with Laemmli buffer. Standard procedures were used for chemiluminescent western blot detection of proteins. Membranes were imaged using a LI-COR Odyssey imager and signals quantitated in the associated ImageStudio software.

### Detergent extraction of potential insoluble protein

We took two approaches to detecting potentially aggregated Aac2^A128P,A137D^. First, spheroplasts were generated using zymolyase followed by lysis in Sample Buffer (containing 8 M Urea + 5% SDS) with or without 2% Sarkosyl. Second, the insoluble material from our typical lysis procedure (see above) was treated with different detergents. For formic acid, pellet was resuspended in 100% formic acid, incubated at 37°C for 70 minutes, followed by SpeedVac drying of the sample and resuspension in Sample Buffer for SDS-PAGE. For guanidine dissociation, pellet was resuspended in 8 M Guanidine HCl, incubated at 25°C for 70 minutes, then sodium deoxycholate was added to a final concentration of 1%. Protein was then precipitated with TCA, washed with acetone at −20°C, and resuspended in Sample Buffer for SDS-PAGE.

### Affinity chromatography

His-tagged Aac2 affinity purification was performed on 1 mg mitochondria per replicate. Yeast mitochondria were isolated as previously described (87). Mitochondria were lysed in “Lysis Buffer” (10% glycerol, 1.5% digitonin, 50 mM potassium acetate, 2 mM PMSF and a protease inhibitor cocktail) including 100 mM NaCl (“low salt”) or 200 mM NaCl (“high salt”) and incubated on ice for 30 minutes. Lysate was centrifuged at 21,000 g for 30 minutes. Imidazole was added to supernatant for a final concentration of 4 mM, which was applied to preequilibrated Ni-NTA agarose beads (Qiagen) and rotated at 4°C for 2 hours. Beads were then washed three times with “Lysis Buffer” (including the corresponding NaCl) but with 0.1% digitonin. Protein was eluted from the beads with 300 mM imidazole, 2% SDS, 10% glycerol in 20 mM HEPES-KOH (pH 7.4). For mass spectrometry, samples were briefly run into an SDS-PAGE gel. Whole lanes were excised, and protein was subject to in-gel trypsin digestion. To ensure reproducibility, the eluate used for mass spectrometry was derived from two independent mitochondrial preparations per strain, and affinity chromatography was performed on two separate days.

HA-tagged Ant1 affinity purification was performed on whole-cell lysate. Each replicate was from an independent transfection of a 10-cm dish of HeLa cells. Briefly, 24 hours after transfection, cells were collected, washed twice in cold PBS and then lysed in 0.5 ml lysis buffer (50 mM HEPES-KOH pH 7.4, 10% glycerol, 100 mM NaCl, 1% Digitonin, 1 mM PMSF, and 1X HALT Protease and Phosphatase Inhibitor Cocktail (Thermo Fisher)) for 30 minutes on ice. Lysate was cleared for 30 minutes at 16,000 g, and supernatant applied to pre-equilibrated Pierce^TM^ anti-HA agarose beads (Thermo Fisher) and incubated overnight with gentle agitation. Beads were subsequently washed 5 times with lysis buffer, except containing 0.1% digitonin, followed by elution with 6% SDS and 10% glycerol, in 50 mM HEPES-KOH pH 7.4.

### Sample processing for mass spectrometry

The excised lanes of yeast eluate (see above) were subjected to in-gel trypsin digestion (88). Briefly, gel pieces were washed with 50 mM ammonium bicarbonate (Acros) in 50% acetonitrile (Fisher), reduced with dithiothreitol (Acros) and alkylated with iodoacetamide (Sigma), washed again, and impregnated with 75 µL of 5 ng/µL trypsin (trypsin gold; Promega) solution overnight at 37°C. The resulting peptides were extracted using solutions of 50% and 80% acetonitrile (ACN) with 0.5% formic acid (Millipore), and the recovered solution dried down in a vacuum concentrator. Dried peptides were dissolved in 60 µL of 0.1% trifluoroacetic acid (TFA, Sigma), and desalted using 2-core MCX stage tips (3M) (89). The stage tips were activated with ACN followed by 3% ACN with 0.1% TFA. Next, samples were applied, followed by two washes with 3% ACN with 0.1% TFA, and one wash with 65% ACN with 0.1% TFA. Peptides were eluted with 75 µL of 65% ACN with 5% NH_4_OH (Sigma), and dried.

Cytosolic fractions from HeLa cells were processed using the FASP method (90). Briefly, in-solution proteins were reduced and denatured with DTT and SDS, mixed with urea to 8 M, and concentrated on a 10 kDa MWCO membrane filter (Pall, OD010C34). Cysteine residues were alkylated using iodoacetamide (Sigma) at room temperature in a dark location for 25 min. The proteins were rinsed with urea and ammonium bicarbonate solutions and digested overnight at 37°C using trypsin gold (Promega) at a ratio of 1:100. The resulting peptides were recovered from the filtrate and a 10 µg aliquot was desalted on 2-core MCX stage tips as above.

### LC-MS methods

Samples were dissolved in 20 to 35 µL of water containing 2% ACN and 0.5% formic acid to 0.25 µg/µL. Two µL (0.5 µg) were injected onto a pulled tip nano-LC column with 75 µm inner diameter packed to 25 cm with 3 µm, 120 Å, C18AQ particles (Dr. Maisch). The column was maintained at 45°C with a column oven (Sonation GMBH). The peptides were separated using a 60-minute gradient from 3 – 28% ACN over 60 min, followed by a 7 min ramp to 85% ACN. The column was connected inline with an Orbitrap Lumos via a nanoelectrospray source operating at 2.2 kV. The mass spectrometer was operated in data-dependent top speed mode with a cycle time of 2.5s. MS^1^ scans were collected at 60000 resolution with AGC target of 6.0E5 and maximum injection time of 50 ms. HCD fragmentation was used followed by MS^2^ scans in the Orbitrap at 15000 resolution with AGC target 1.0E4 and 100 ms maximum injection time.

### Database searching and label-free quantification

The MS data was searched using SequestHT in Proteome Discoverer (version 2.4, Thermo Scientific) against the *S. Cerevisiae* proteome from Uniprot, containing 6637 sequences, concatenated with common laboratory contaminant proteins. Enzyme specificity for trypsin was set to semi-tryptic with up to 2 missed cleavages. Precursor and product ion mass tolerances were 10 ppm and 0.6 Da, respectively. Cysteine carbamidomethylation was set as a fixed modification. Methionine oxidation was set as a variable modification. The output was filtered using the Percolator algorithm with strict FDR set to 0.01. Label-free quantification was performed in Proteome Discoverer with normalization set to total peptide amount.

### Label-free quantitation data processing

For analysis, label-free quantitation with Proteome Discoverer software generated protein abundances for each sample. Protein abundances were then analyzed using Metaboanalyst software (91). First, we refined the protein list to include only proteins whose fold-change is > 1.5 (*p* < 0.1) in Aac2-HIS6 eluate compared with the null control. Then, to eliminate any bias introduced by mutant bait protein Aac2^A128P^-HIS6 being present at ∼50% the level of Aac2-HIS6 (see Fig. S3C & S4A), we normalized each prey protein abundance by that sample’s bait protein level (i.e. Aac2 level). Finally, we performed multiple *t* testing on these values, which generated the FDR-corrected *p*-values and were ultimately normalized to wild-type level for presentation in Fig. 4C, S3I, S4B and S4E. Gene Ontology and other enrichment analyses were performed using STRING version 11.0 (92).

For the analysis shown in Fig. 5F, the protein list was manually curated to include only proteins that had a Gene Ontology Cellular Component term for “mitochondrion”, and not that of any other organelle. Proteins with missing values were excluded. The levels of each protein were averaged within an experimental group and those average values were normalized to the average value for wild-type Ant1-transfected samples. It is these wild-type-normalized average values for each protein that were subject to student’s *t* test to probe for a significant difference in mitochondrial protein levels present in the cytosol of mutant compared with wild-type Ant1-transfected HeLa cells.

### Targeted quantitative proteomics

Unused yeast peptides from the label-free quantification experiment were used for absolute quantification using parallel reaction monitoring (PRM) on the instrument described above. The instrument method consisted of one MS scan at 60000 resolution followed by eight targeted Orbitrap MS^2^ scans at 30000 resolution using quadrupole isolation at 1.6 m/z and HCD at 35%.

Two heavy-labeled proteotypic peptides for Aac2 and Tim22 were purchased from New England Peptide. Their sequences were TATQEGVISFWR and SDGVAGLYR; and VYTGFGLEQISPAQK and TVQQISDLPFR respectively, each with N-terminal ^13^C_6_ ^15^N_4_ arginine or ^13^C_6_ ^15^N_2_ lysine. An equal portion of the peptides contained in each gel band was combined with a mixture of proteotypic peptides resulting in an on-column load of 1 fmol of each Tim22 heavy peptide and 250 fmol of each Aac2 heavy peptide. Assays were developed for the doubly-charged precursor and the 4 or 5 most intense singly-charged product y-ions. The data was analyzed in Skyline (version 20.2) (93).

### qRT-PCR

Total RNA was isolated from ∼1.5 X 10^7^ yeast cells using Quick-RNA Fungal/Bacterial Microprep Kit (Zymo Research). Quantitative real-time PCR (qRT-PCR) was executed using 50 ng RNA in Power SYBR Green RNA-to-Ct 1-step Kit (Applied Biosystems). A CFX384 Touch Real-Time PCR Detection System (BioRad) was used with the following cycling parameters: 48°C 30 minutes, 95°C 10 minutes, followed by 44 cycles of 95°C 15 seconds, 60°C 1 minute. *TFC1* was used as reference, as its mRNA level was previously established as stable across culture conditions (94). The specificity of each primer pair was validated using RNA extracted from knockout strains. Ct values were determined using CFX Maestro software (BioRad) and analyzed manually using the 2^−ΔΔCT^ method.

### *in vitro* mitochondrial protein import assays

The *AAC2* variants were cloned into pGEM4z plasmids. The constructs were used for coupled *in vitro* transcription and translation, using a cell free system based on reticulocyte lysate. The wild-type strains YPH499 and BY4741 and yeast expressing Tom40-HA were grown in YPG (1% [w/v] yeast extract, 2% [w/v] bacto-peptone and 3% [v/v] glycerol) at 30°C. Mitochondria were isolated by differential centrifugation. The import of the Aac2 variants were performed as described (18). Isolated mitochondria were incubated for different time periods with radiolabeled Aac2 variants in the presence of 4 mM ATP and 4 mM NADH in import buffer (3% [w/v] bovine serum albumin; 250 mM sucrose; 80 mM KCl; 5 mM MgCl_2_; 5 mM methionine; 2 mM KH_2_PO_4_; 10 mM MOPS/KOH, pH 7.2). The import reaction was stopped by addition of 8 µM antimycin A, 1 µM valinomycin and 20 µM oligomycin (AVO; final concentrations). In control reactions, the membrane potential was depleted by addition of the AVO mix. Subsequently, mitochondria were washed with SEM buffer (250 mM sucrose; 1 mM EDTA; 10 mM MOPS-KOH, pH 7.2), lysed with 1% (w/v) digitonin in lysis buffer (0.1 mM EDTA; 50 mM NaCl; 10% [v/v] glycerol; 20 mM Tris-HCl, pH 7.4) for 15 min on ice analysis and protein complexes were separated on blue native gels. To remove non-imported preproteins, mitochondria were treated with 50 µg/ml proteinase K for 15 min on ice. The protease was inactivated by addition of 1mM phenylmethylsulfonyl fluoride (PMSF) for 10 min on ice. To study the accumulation of AAC2 variants at the TOM translocase, the AAC2 variants were imported into Tom40-HA mitochondria followed by affinity purification via anti-HA beads (Roche) (18). After the import reaction, mitochondria were lysed with 1% (w/v) digitonin in lysis buffer for 15 min on ice. After removing insoluble material, the mitochondrial lysate was incubated with anti-HA matrix for 60 min at 4°C. Subsequently, beads were washed with an excess amount of 0.1% (w/v) digitonin in lysis buffer and bound proteins were eluted under denaturing conditions using SDS sample buffer (2% [w/v] SDS; 10% [v/v] glycerol; 0.01% [w/v] bromophenol blue; 0.2% [v/v] β-mercaptoethanol; 60 mM Tris/HCl, pH 6.8). Precursors of Aac2 and its variants were arrested at different import stages by depleting ATP and/or the Δψ in the import reaction as described (18, 45).

### Pulse-chase analysis of Aac2^A128P, A137D^

Cells with disrupted endogenous *AAC2* but expressing *GAL10*-*aac2^A128P, A137D^*, with or without the disruption of *YME1* and *PDR5*, were grown in YPD at 30°C overnight. Cells were then subcultured in complete galactose plus raffinose medium for 4 hours at 30°C. Cycloheximide was added to a concentration of 1 mg/ml, with or without the addition of 100 μM MG132. Cycloheximide chase was pursued at 30°C for 30, 60 and 120 min before cells were collected for western blot analysis.

### Expression of *ANT1* in HeLa cells

HeLa cells were cultured in DMEM (Gibco) with 10% fetal bovine serum (Sigma) at 37°C in a humidified atmosphere of 5% CO_2_. *ANT1* cDNA was cloned into pCDNA3.1 with an HA epitope added to the C-terminus, as previously described (65). Mutant *Ant1* alleles were generated by *in vitro* mutagenesis using QuikChange Site-Directed Mutagenesis (Stratagene) and confirmed by sequencing. Cells were transfected using Lipofectamine 3000 (Invitrogen) according to the manufacturer’s protocol and harvested 24 hours after transfection for all experiments.

### HeLa cell fractionation

One 10-cm dish per sample was harvested, washed twice in cold PBS, homogenized in 1 mL Isotonic Buffer (250 mM sucrose, 1 mM EDTA, 10 mM Tris-HCl, pH 7.4,) including HALT Protease and Phosphatase Inhibitor Cocktail (Thermo Fisher) with 10 slow strokes in a 2 ml Dounce homogenizer (KIMBLE, pestle B, clearance = 0.0005-0.0025 inches). Homogenate was centrifuged at 600 g for 15 minutes. To pellet mitochondria, supernatant was spun at 10,000 g for 25 minutes. Supernatant was the cytosolic fraction and pellet (the mitochondrial fraction) was used subsequently for protease protection or western blotting. All steps were performed on ice.

### Protease protection assay in HeLa cells

For protease protection, isolated mitochondria from four 10-cm plates were combined to generate each replicate indicated in Fig. 5E. All replicates were from independent transfections on different days. 20 μg aliquots of mitochondria were pelleted and resuspended in either isotonic buffer (as above, without protease inhibitors) or hypotonic buffer (10 mM KCl, 2 mM HEPES, pH 7.2) for swelling on ice for 20 minutes. Where indicated, proteinase K was added at a final concentration of 7 μg/ml, incubated at room temperature for 20 minutes, and inhibited with 5 mM PMSF on ice for 15 minutes. Where indicated, 1% Triton X-100 treatment on ice for 10 minutes was used to lyse the mitochondrial membranes. Ultimately, half of each reaction was loaded onto two independent gels for western blotting. Immunoblotting of reference proteins in Fig. 5D was done for each replicate. If a reference protein was not properly protected/degraded, this sample was discarded.

### Apoptosis assay and relative Δψ determination

Cells were harvested 24 hours post-transfection and processed for flow cytometry detection of apoptosis and membrane potential after staining with propidium iodide and annexin V-FITC and JC-1 respectively (65).

### *Ant1^A114P,A123D^*/+ knock-in mice

All procedures were approved by the Animal Care and Use Committee (IACUC) at State University of New York Upstate Medical University and were in accordance with guidelines established by the National Institutes of Health.

The *Ant1* targeting vector was prepared by recombineering as previously described (95). Briefly, 13.8 kb of *Ant1* genomic sequence containing all 4 exons plus 4.4 kb of 5’ upstream and 4.96 kb 3’ downstream untranslated sequences was retrieved from the RP24-108A1 BAC clone obtained from the BACPAC Resources Center (Children’s Hospital Oakland Research Institute, Oakland, CA). The first loxP site together with the Frt-PGKnew-Frt cassette was inserted approximately 0.4 kb 3’ of exon 4, which contains the polyA signal sequence and the 3’UTR. Two unique restriction sites, *Asc* I and *Asi*SI, were also introduced into the 3’ end of the second loxP site to allow insertion of the *Ant1* mini-cDNA containing the A114P and A123D knock-in mutations as well as the His-tag at the carboxyl terminus.

The *Ant1* mini-cDNA was prepared by fusion PCR using primers with sequences overlapping different exons. The mini-cDNA contains *Asc* I and *Asi* SI unique restriction sites in the 5’ and 3’ end, respectively, together with 254 bp of intron 1 sequence together with the splice acceptor followed by exon 2 with the two mutations, exon 2 and 4 and 70 bp of 3’ downstream sequence. The mini cDNA was cloned into pSK+ and sequenced to confirm its identity prior to insertion into the *Asc* I and *Asi* SI sites in the targeting vector. The final targeting vector was then linearized by *Not* I digestion, purified, and resuspended in PBS at 1 *m*g/ul for electroporation into ES cells derived from F1 (129Sv/C57BL6j) blastocyst. Targeted ES clones were identified by long range nested PCR using Platinum HiFi Taq (Invitrogen).

Chimeric animals were generated by aggregation of ES cells with the CD1 morula. Chimeric males were bred with ROSA26-Flpe female (Jackson Labs stock no: 009086) to remove the PGKnew cassette and generate F1 pups with *Ant1* floxed allele. Positive pups were identified by PCR genotyping using primer Lox gtF (5’-ATCCATCTCAAAGGCAAACG-3’) and Lox gtR (5’-AAATTCCCTGCAGGCTTATG-3’) to detect a fragment of 364 bp specific to the 5’-Lox site and a fragment of 270 bp specific to the wild-type allele. A heterozygous floxed male was bred with Hprt-Cre female (Jackson Labs stock no: 004032). The same primers were used for genotyping knock-in mice, with the 270 bp band present only with the knock-in allele, and no band produced from a wild-type locus with lacking loxP sites. The mixed background heterozygous knock-in mice (i.e. Ant1^A114P,A123D^/+) male mice were backcrossed with C57BL/6NTac females (Taconic Catalog no: B6-F) for 7 generations before experiments were performed.

### Mouse Histology

For spinal cord histology, mice were sacrificed by isoflurane overdose, followed by intracardial perfusion with PBS followed by PBS + 4% paraformaldehyde (PFA). Perfused mice were soaked in PBS + 4% PFA overnight at 4°C, and then spinal cord was dissected and cryopreserved with increasing concentrations of sucrose in PBS. Tissue was then embedded in OCT and snap-frozen in 2-methylbutane on liquid N_2_. Tissue was sectioned at 8 μm with a cryostat (Leica) and sections stained with Cresyl Violet Stain Solution according to the manufacturer’s protocol (Abcam) or used for indirect immunofluorescence. For the latter, tissue was blocked with 10% horse serum and then incubated with rat monoclonal anti-GFAP antibody (Invitrogen) overnight at 4°C. After rinsing the tissue was incubated with fluorescently-conjugated anti-rat secondary antibody (Immunoresearch West Grove, PA).

For muscle histology, the soleus muscles were quickly dissected and fresh-frozen in 2-methylbutane on liquid N_2_. Tissue was cryosectioned at 10 μm and stained with hematoxylin and eosin (H&E) using standard procedures. Feret’s diameter of myofibers was determined using ImageJ software; all soleus myofibers from a single muscle section per mouse (n > 340 fibers per mouse) were quantitated by a blinded observer. For SDH only staining, sections were air-dried, incubated at 37°C for 45 minutes in SDH medium (0.1 M succinic acid, 0.1 M sodium phosphate buffer pH 7, 0.2 mM phenazine methosulfate, + 1 mg/mL NBT added fresh), drained, fixed in 10% formalin, rinsed well with water and mounted in water soluble mounting medium. For sequential COX/SDH staining, sections were air-dried, incubated in Cytochrome c Medium (0.5 mg/mL 3’3’-diaminobenzidine tetrahydrochloride (DAB), 75 mg/mL sucrose in 50 mM sodium phosphate buffer, pH 7.4, with freshly added cytochrome c and catalase at 1 mg/ml and 0.1 mg/ml respectively) at 37°C for 1 hour, followed by a quick wash in water and SDH staining as described above. Abnormal COX/SDH-stained fibers were scored manually from decoded images of whole soleus sections.

### Electron Microscopy

For electron microscopy, Ant1^A114P,A123D^/+ mice and littermate controls were processed as previously described (96). Briefly, mice were anesthetized with isoflurane and perfused intracardially with PBS initially, followed by fixative (1% paraformaldehyde, 1% glutaraldehyde, 0.12 M sodium cacodylate buffer pH 7.1, and 1 mM CaCl_2_). Perfused animals were refrigerated overnight, and CNS tissues were dissected the next day and processed for TEM. The samples were examined with a JOEL JEM1400 transmission electron microscope and images were acquired with a Gaten DAT-832 Orius camera.

### Bioenergetic analysis

For mitochondrial respiration experiments, the mice were sacrificed by decapitation via guillotine without the use of CO_2_ asphyxiation or anesthetic. Skeletal muscle mitochondria were isolated and respiration measured as previously described (97). Briefly, after mitochondria isolation by differential centrifugation, oxygen tension was measured using an Oxygraph Plus oxygen electrode (Hansatech Instruments) in 0.5 ml Experimental Buffer containing 150 μg mitochondria in a temperature-controlled 37°C chamber. For complex I measurements, glutamate and malate were added for a final concentration of 5 mM and 2.5 mM, respectively. For complex II measurements, rotenone was added before succinate, for final concentrations of 5 μM and 10 mM respectively.

### Protease protection assay of mouse skeletal muscle mitochondria

For protease protection assay, skeletal muscle mitochondria were isolated as for bioenergetic analysis with slight modifications, including the addition of 1 mM PMSF in the homogenization buffer, and resuspending mitochondria in a modified isotonic buffer lacking BSA (75 mM sucrose, 215 mM mannitol, 1 mM EGTA in 20 mM HEPES-KOH pH 7.4). 80 μg aliquots were treated with the indicated concentrations of proteinase K (Sigma) for 30 minutes at room temperature and quenched with 5 mM PMSF on ice for 10 minutes. Where indicated, mitochondria were lysed with 1% Triton X-100 for 30 minutes on ice. Laemmli buffer was ultimately added to the samples such that the final protein concentration was ∼40 μg mitochondria per 15 μl, which was the amount loaded onto the SDS-PAGE gel for Western blotting. Such high protein amounts were required for detection of Ant1^A114P,A123D^.

### Mouse skeletal muscle sub-cellular fractionation

Five biological replicates of wild-type and Ant1^A114P,A123D^/+ muscle samples were fractionated to obtain cytosolic and mitochondrial fractions using differential centrifugation, as follows. 100 mg Quadriceps muscle was rapidly dissected and immediately placed in ice cold PBS. Muscle was minced, centrifuged for 1 min at 500g, and resuspended in 1 ml buffer STM (250 mM sucrose, 50 mM Tris-HCl pH 7.4, 5 mM MgCl2 plus HALT protease and phosphatase inhibitors added fresh). Tissue was homogenized in a 2 ml Dounce homogenizer (0.15-0.25 mm clearance) with two strokes using a bench top drill press set to 570 rotations per minute. Homogenate was centrifuged for 15 minutes at 800g twice, and the pellets were discarded after both spins. To obtain the mitochondrial fraction, supernatant was centrifuged for 11,000g for 10 minutes. Mitochondrial fraction was washed twice in STM and frozen for further analysis. Meanwhile, the supernatant was centrifuged for 30 minutes at 21,000g twice to remove any contaminating mitochondria from the cytosolic fraction.

### Mouse skeletal muscle cytosolic protein digestion, labeling, cleanup, and fractionation

Skeletal muscle cytosolic fractions were then processed for multiplexed quantitative mass spectrometry as follows. Samples were buffer exchanged on a 3 kDa molecular weight cutoff filter (Amicon 3k Ultracel) using 4 additions of 50 mM triethylammonium bicarbonate, pH 8.0 (Thermo). Following a Bradford assay, 50 µg of each cytosolic fraction was taken for digestion using an EasyPep Mini MS sample prep kit (Thermo, A40006). To each buffer-exchanged sample, 70 µL of lysis buffer was added followed by 50 µL of reduction solution and 50 µL of alkylating solution. Samples were incubated at 95°C for 10 minutes, then cooled to room temperature. To each sample 2.5 µg of trypsin / Lys-C protease was added and the reaction was incubated at 37°C overnight. TMT reagents were reconstituted with 40 µL acetonitrile (ACN) and the contents of each label added to a digested sample. After 60 min, 50 µL of quenching solution was added, consisting of 20% formic acid and 5% ammonium hydroxide (v/v) in water. The labeled digests were cleaned up by a solid-phase extraction contained in the EasyPep kit, and dried by speed-vac. The ten cytosolic fractions were dissolved in 50 µL of 30% ACN and 0.1% formic acid (v/v) in water, combined, and dried again.

Following an LC-MS experiment to check digestion and labeling quality of the pooled samples, these were fractionated using a Pierce High pH Reversed-Phase Peptide Fractionation Kit (part # 84868), per the manufacturer’s instructions for TMT-labeled peptides. In brief, samples were dissolved in 300 µL of 0.1% trifluoroacetic acid in water and applied to the conditioned resin. Samples were washed first with water and then with 300 µL of 5% ACN, 0.1% triethylamine (TEA) in water. The second wash was collected for analysis. Peptides were step eluted from the resin using 300 µL of solvent consisting of 5 to 50% ACN with 0.1% TEA in eight steps. All collected fractions were dried in a speed-vac.

### LC-MS/MS for TMT-labeled samples

Dried fractions were reconstituted in 50 µL of load solvent consisting of 3% ACN and 0.5% formic acid in water. Of these, 2 µL were injected onto a pulled tip nano-LC column (New Objective, FS360-75-10-N) with 75 µm inner diameter packed to 25 cm with 2.2 µm, 120 Å, C18AQ particles (Dr. Maisch GmbH). The column was maintained at 50°C with a column oven (Sonation GmbH, PRSO-V2). The peptides were separated using a 135-minute gradient consisting of 3 – 12.5% ACN over 60 min, 12.5 – 28% over 60 min, 28 - 85 % ACN over 7 min, a 3 min hold, and 5 min re-equilibration at 3% ACN. The column was connected inline with an Orbitrap Lumos (Thermo) via a nanoelectrospray source operating at 2.3 kV. The mass spectrometer was operated in data-dependent top speed mode with a cycle time of 3s. MS^1^ scans were collected from 375 – 1500 m/z at 120,000 resolution and a maximum injection time of 50 ms. HCD fragmentation at 40% collision energy was used followed by MS^2^ scans in the Orbitrap at 50,000 resolution with a 105 ms maximum injection time.

### Database searching and reporter ion quantification

The MS data was searched using SequestHT in Proteome Discoverer (version 2.4, Thermo Scientific) against the *M. Musculus* proteome from Uniprot, containing 50887 sequences, concatenated with common laboratory contaminant proteins. Enzyme specificity for trypsin was set to semi-tryptic with up to 2 missed cleavages. Precursor and product ion mass tolerances were 10 ppm and 0.6 Da, respectively. Cysteine carbamidomethylation, TMT 10-plex at any N-terminus and TMT 10-plex at lysine were set as a fixed modifications. Methionine oxidation was set as a variable modification. The output was filtered using the Percolator algorithm with strict FDR set to 0.01. Quantification parameters included the allowance of unique and razor peptides, reporter abundance based on intensity, lot-specific isotopic purity correction factors, normalization based on total peptide amount, protein ratio based on protein abundance, and hypothesis testing (ANOVA) for individual proteins.

### Grip strength measurements

Grip strength measurements were performed using Bio-CIS software connected with the Grip Strength Test Model GT3 according to manufacturer’s protocol for forelimb-only measurement using the grid (BIOSEB). Briefly, mice were held by the tail above the grid that’s connected to a force meter, slowly lowered to allow the forelimbs to grip the grid, and then slowly and smoothly pulled horizontally along the axis of the sensor until the grasp was released. The maximum force generated is recorded by the software and reported as the average of 5 consecutive trials or the maximum force generated over 5 trials.

### Statistical Analysis

Statistical analyses were performed using GraphPad Prism. For details on statistical testing of specific data, please see Figure Legends.

## Acknowledgments

We thank Nikolaus Pfanner for support and anti-sera, Joyce Qi for help with electron microscopy, and Siu-Pok Yee (University of Connecticut) for Ant1^A114P,A123D^ knock-in mouse generation. We’re also grateful to Yumiko Umino and Eduardo Solessio for confirming visual acuity and contrast sensitivity in Ant1^A114P,A123D^/+ mice.

## Author contributions

Conceptualization: LPC, XJC

Methodology: LPC, TB, XJC

Formal Analysis: LPC

Investigation: LPC, XW, JS, EdJ, KS

Visualization: LPC, XJC Supervision: PM, FM, TB, XJC

Writing—original draft: LPC

Writing—review & editing: LPC, JS, PM, TB, XJC

Funding acquisition: LPC, TB, XJC

## Data and materials availability

All software used in this study is publicly available (Resources Table in Methods section), and there was no code generated as part of this study. The small datasets generated as part of this study are either present as a supplemental table or will be made available from the corresponding author on request.

