## Supplemental data for "Mitochondrial protein import clogging as a mechanism of disease"

**This PDF file includes:**

Figs. S1 to S7

**Other Supplementary Materials for this manuscript include the following:**
[use this section only if you have movies, audio or data files]
Movies S1 to S2
Data S1 to S3

**Fig. S1.**

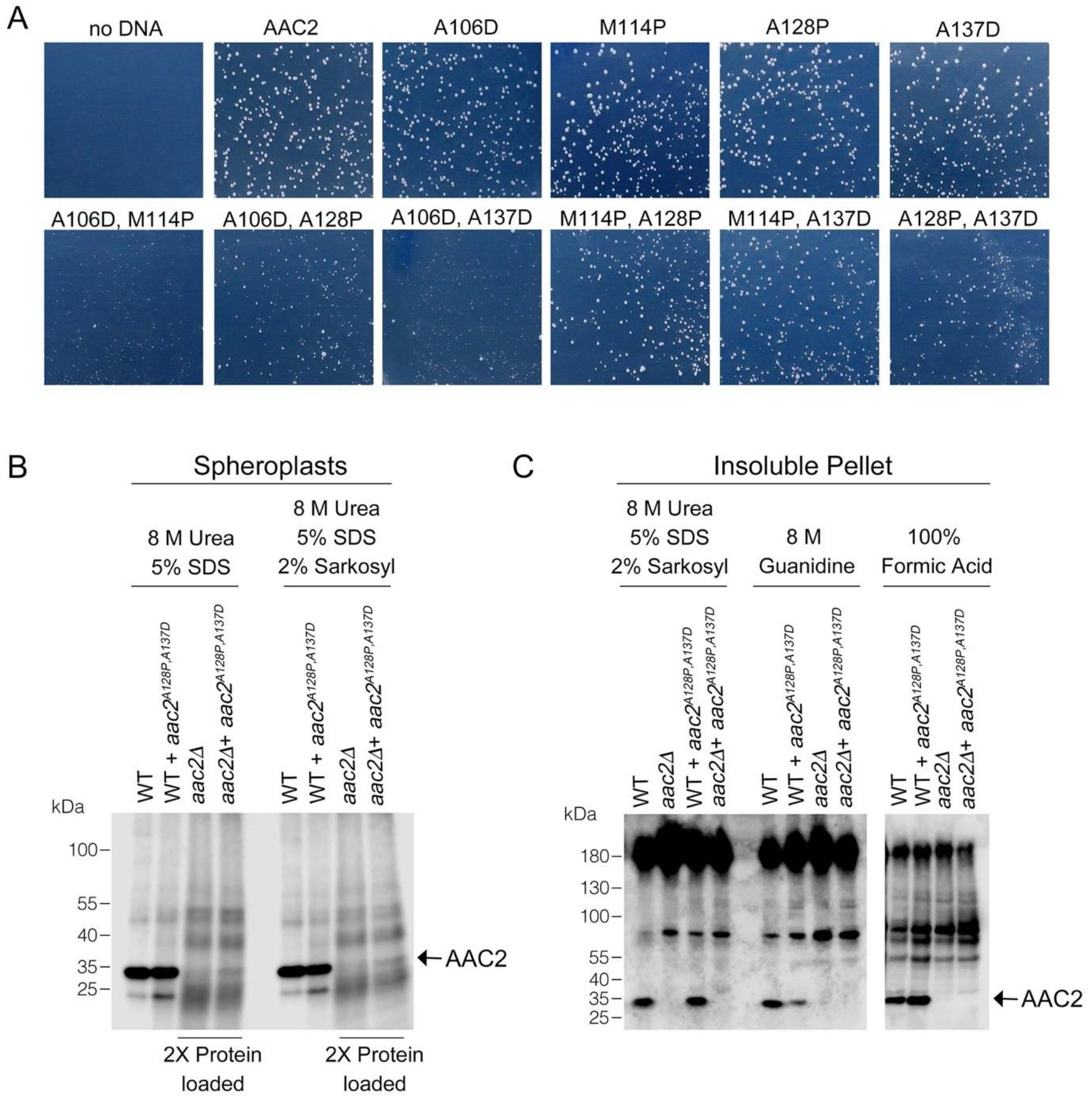


**Fig. S1. Toxicity and low-level accumulation of double mutant Aac2 proteins. (A)** Yeast transformants expressing double mutant alleles of *aac2* form small colonies on selective medium, which indicates the toxicity of input plasmid DNA. The yeast M2915-6A strain was transformed with the centromeric vector pRS416 (*URA3*) expressing wild-type or mutant *aac2* alleles and transformants were grown on minimal glucose medium lacking uracil at 30°C for 3 days. This serves as a control for Figure 1B. **(B)** Immunoblot analysis of Aac2 after protein extraction from spheroplasts using the detergents indicated. **(C)** Immunoblot analysis of Aac2 in protein extracted from the detergent-insoluble pellet with additional solubilization conditions after a first round of extraction using 8 M urea plus 5% SDS.

**Fig. S2.**

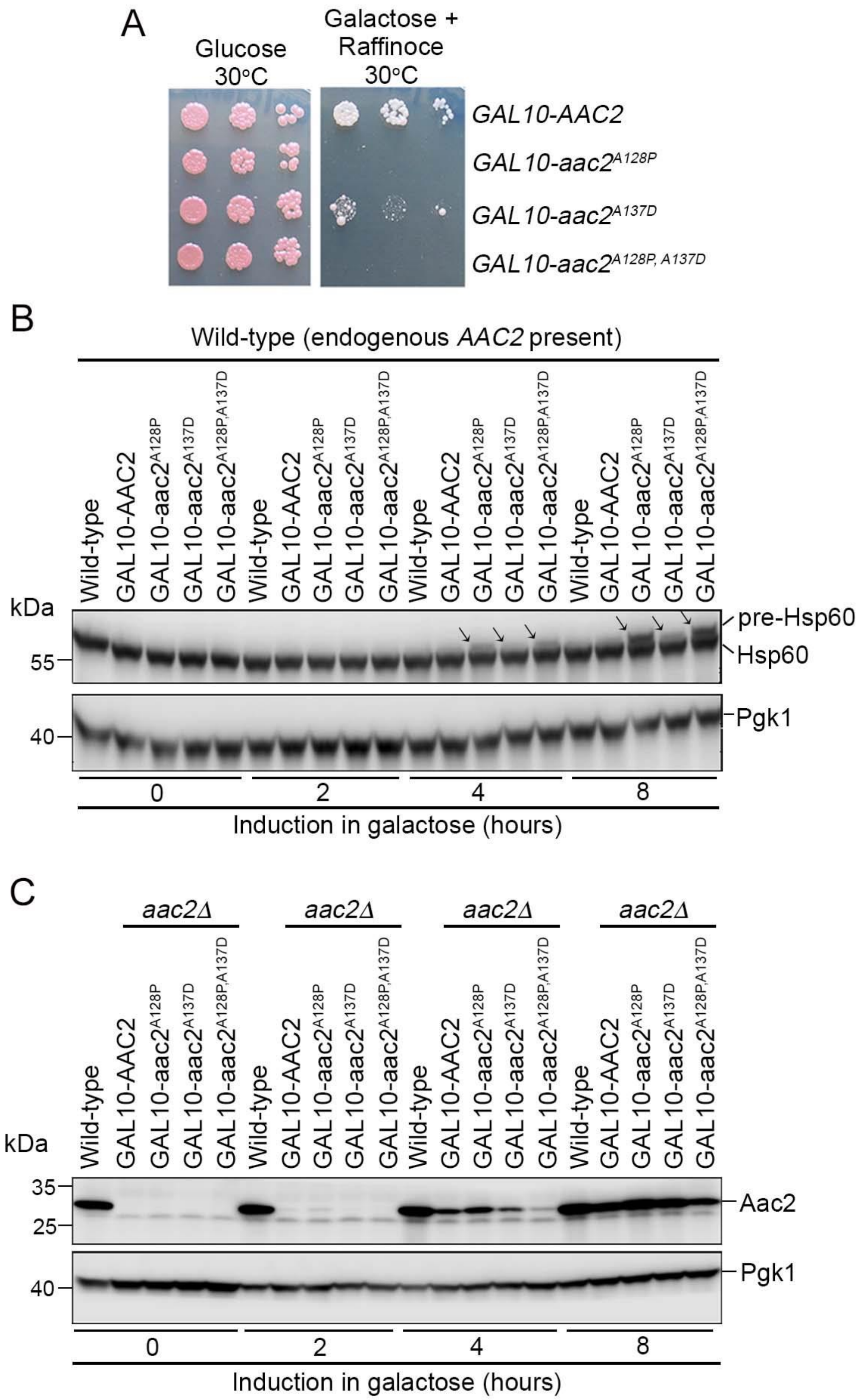


**Fig. S2. Acute expression of mutant Aac2 is toxic and impairs protein import.**

**(A)** Expression of chromosomally integrated *aac2*$^{A128P}$, *aac2*$^{A137D}$ and *aac2*$^{A128P, A137D}$ from the

*GAL10* promoter on complete galactose plus raffinose medium inhibits cell growth. **(B)** Time course study showing that acute expression of the chromosomally integrated *aac2*$^{A128P}$, *aac2*$^{A137D}$ and *aac2*$^{A128P, A137D}$ from the *GAL10* promoter leads to the accumulation of Hsp60 precursor in the presence of a wild-type copy of *AAC2*. **(C)** Western-blot showing expression of chromosomally integrated *aac2*$^{A128P}$, *aac2*$^{A137D}$ and *aac2*$^{A128P, A137D}$ from the *GAL10* promoter in an *aac2Δ* strain background.

**Fig. S3.**

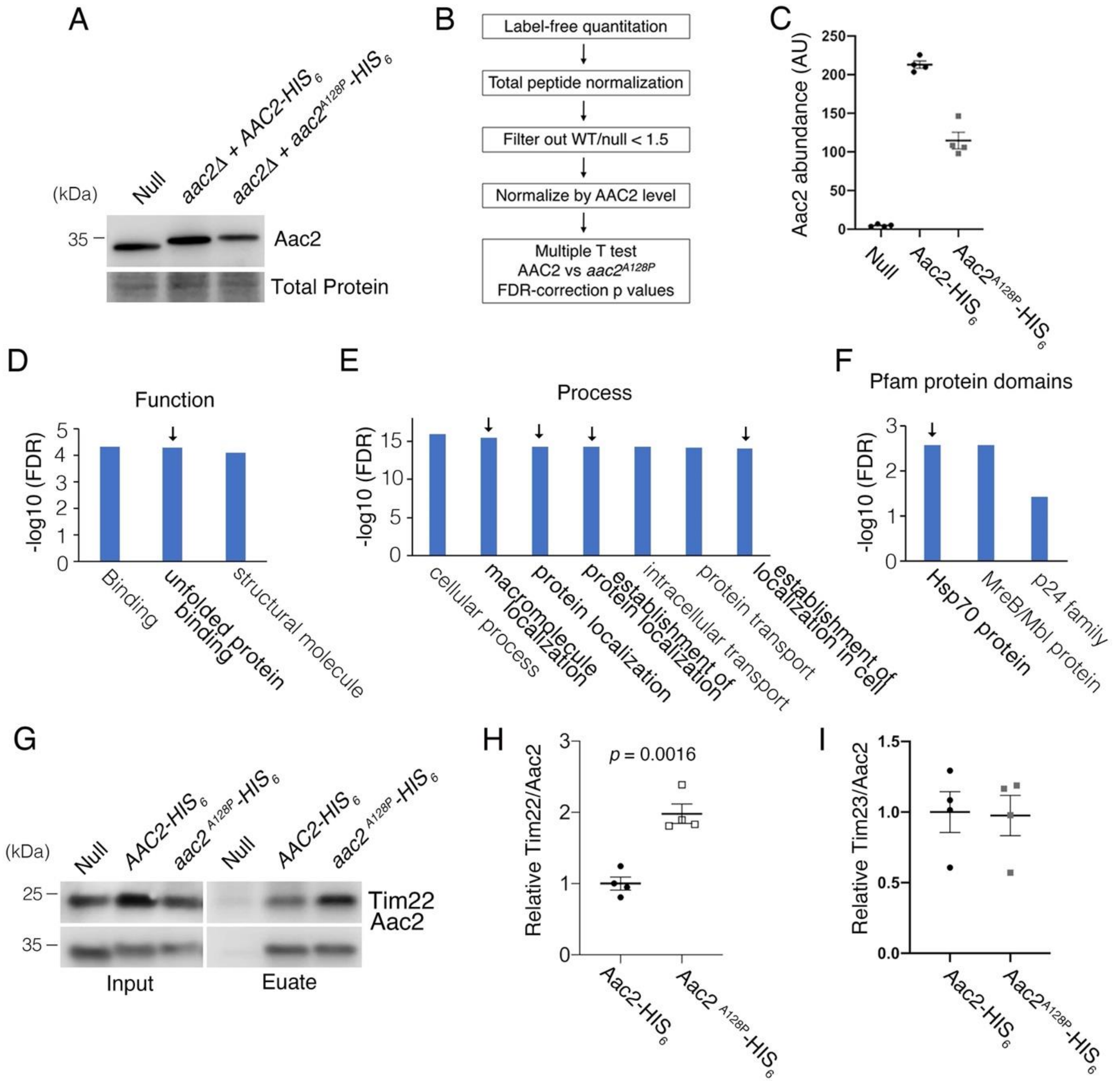


**Fig. S3. Affinity purification in low salt conditions suggested that Aac2$^{A128P}$ accumulates along the carrier protein import pathway. (A)** Immunoblot analysis validating the HIS$_6$-tagged Aac2 proteins. **(B)** Flow chart of label-free quantitative mass spectrometry data processing strategy. See Methods for further details. **(C)** Label-free quantitative mass spectrometry demonstrated that Aac2$^{A128P}$-HIS$_6$ was about half as abundant as Aac2-HIS$_6$ after total peptide normalization. This is consistent with immunoblot in (A), and underscores the need to normalize prey protein levels by bait (i.e. Aac2) protein levels. These data are from the low-salt experiment. **(D)** Three most significant Gene Ontology (GO) Molecular Function terms overrepresented among proteins that are significantly enriched in Aac2$^{A128P}$-$HIS_6$ eluate in the low-salt experiment. **(E)** Seven most significant GO Biological Process terms overrepresented among proteins that are significantly enriched in Aac2$^{A128P}$-$HIS_6$ eluate in the low-salt experiment. **(F)** Three most significant protein families overrepresented among proteins that are significantly enriched in Aac2$^{A128P}$-$HIS_6$ eluate in the low-salt experiment, as determined from the Pfam database. **(G)** Immunoblot analysis validating preferential co-purification of Tim22 with Aac2$^{A128P}$-$HIS_6$ compared with Aac2-$HIS_6$. "Null" strain is wild-type lacking any $HIS_6$-tagged proteins. **(H)** Quantitation from four independent affinity purifications followed by immunoblotting, as shown in (D). *P* value was calculated with student's *t* test. **(I)** The association of Aac2$^{A128P}$ with Tim23, a known binding partner, was not increased compared with wild-type in the low-salt experiment. These data are from label-free quantitative mass spectrometry of pull-down products.

**Fig. S4.**

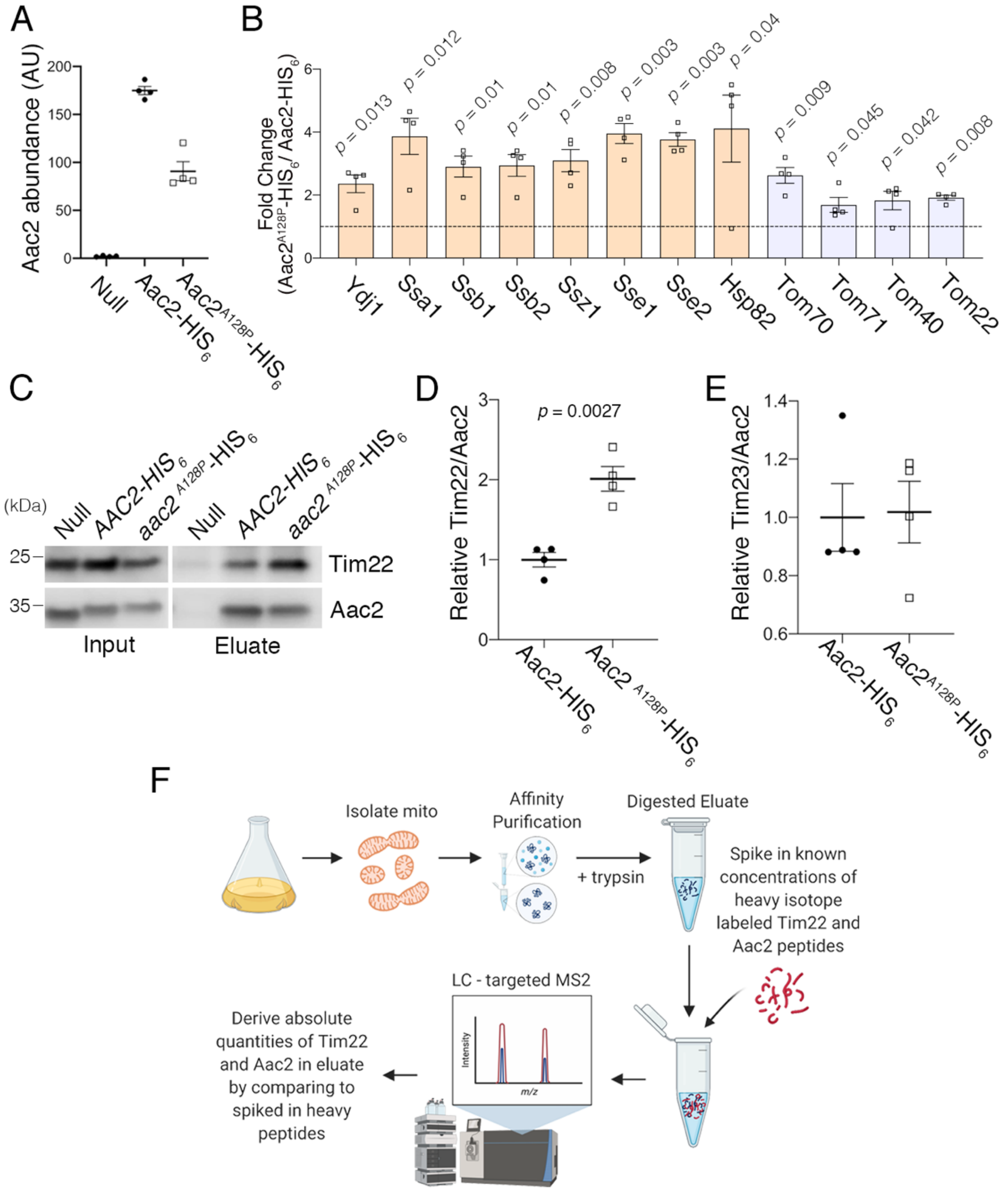


**Fig. S4. Affinity purification in high salt conditions confirmed that Aac2$^{A128P}$ accumulates along the carrier protein import pathway. (A)** Label-free quantitative mass spectrometry demonstrated that Aac2$^{A128P}$-HIS$_6$ was about half as abundant as Aac2-HIS$_6$ after total peptide normalization. This is consistent with the low salt experiment (Figure S3C) and underscores the need to normalize prey protein levels by bait (i.e. Aac2) protein levels. **(B)** Co-purified proteins significantly enriched in $Aac2^{A128P}$-$HIS_6$ eluate compared with Aac2-$HIS_6$ under high-salt purification conditions. FDR-corrected *p* values calculated by multiple *t* test analysis. See Methods for details on abundance values. **(C)** Immunoblot analysis indicating preferential co-purification of Tim22 with $Aac2^{A128P}$-$HIS_6$ compared with Aac2-$HIS_6$ in high-salt conditions. **(D)** Quantitation from four independent affinity purifications followed by immunoblotting, as in (C). *P* value was calculated with student's *t* test. **(E)** The association of $Aac2^{A128P}$ with Tim23, a known binding partner, was not increased compared with wild-type in the high salt experiment. **(F)** Schematic of the parallel reaction monitoring approach for targeted proteomic quantification of Aac2 and Tim22 in pull-down products.

**Fig. S5.**

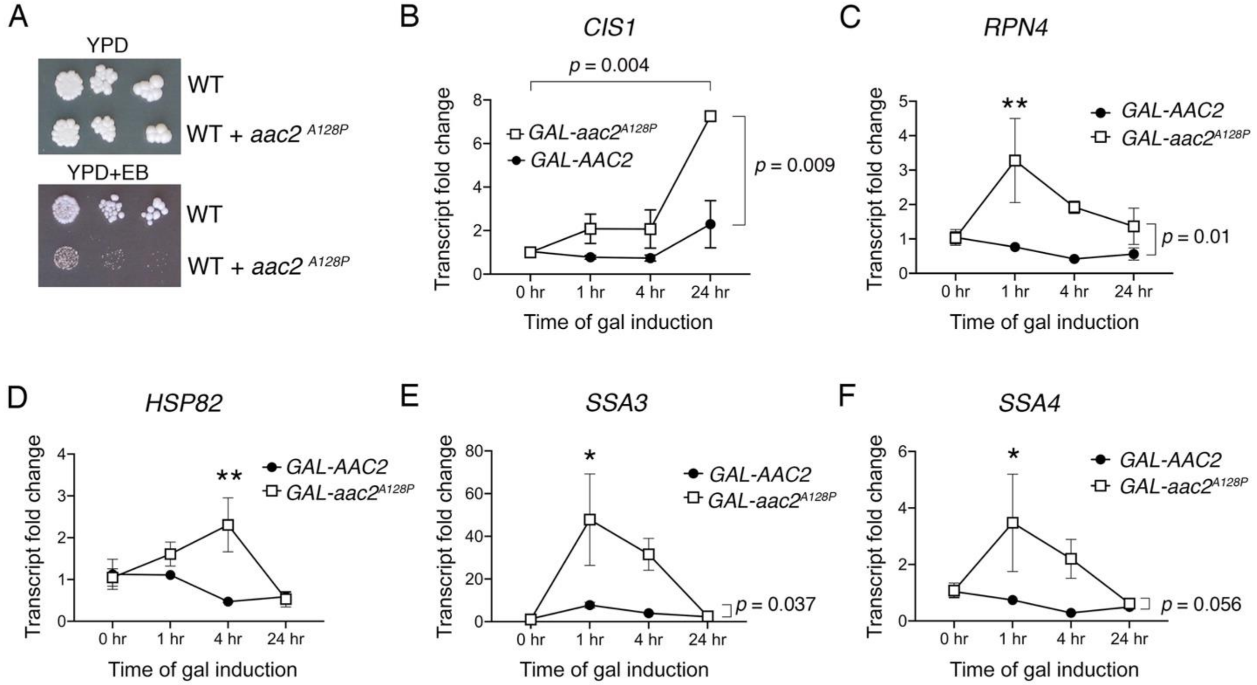


**Fig. S5. Cellular responses to *aac2*$^{A128P}$ expression support mitochondrial protein import clogging. (A)** *aac2*$^{A128P}$ expression is not compatible with elimination of mitochondrial DNA via growth on ethidium bromide (EB) medium. Cells were grown at 30$^{o}$C for 3-4 days. **(B – F)** qRT-PCR analysis monitoring the expression of *CIS1*, *RPN4*, *HSP82*, *SSA3* and *SSA4* after galactose-induced expression of *AAC2* or *aac2*$^{A128P}$. Three biological and two technical replicates were performed for each sample. *TFC1* levels used as reference. *P* values calculated with a two-way repeated measures ANOVA followed by Sidak's multiple comparisons test.

**Fig. S6.**

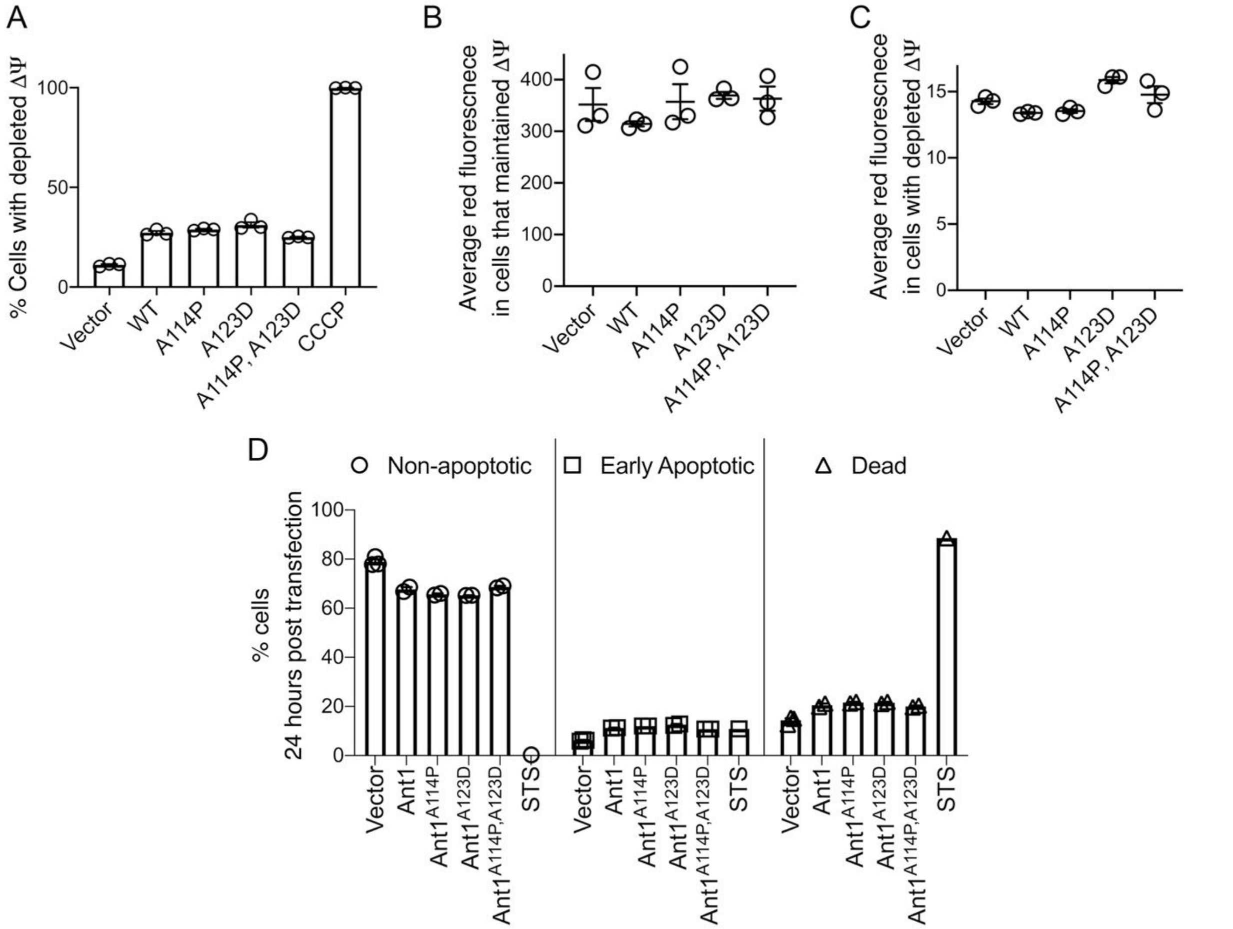


**Fig. S6. Mutant Ant1 does not reduce Δψ or increase apoptosis in HeLa cells. (A)** Mutant Ant1 expression does not increase the fraction of cells with depleted Δψ compared with wild-type Ant1, as indicated by flow cytometry analysis after JC-1 dye staining. Cells treated with the ionophore Carbonyl cyanide m-chlorophenyl hydrazone (CCCP) served as control for Δψ depletion. **(B)** Among cells that maintained Δψ, neither wild-type nor mutant Ant1 transfection reduced the average red fluorescence after JC-1 staining and flow cytometry analysis. The JC-1 dye fluoresces red when it aggregates at high concentration, i.e. after it's Δψ-dependent uptake by mitochondria. In the monomer form in the cytosol, it fluoresces green. **(C)**

Among cells with depleted $\Delta\psi$, neither wild-type nor mutant Ant1 reduced the average red fluorescence after JC-1 staining and flow cytometry analysis. **(D)** Transient expression of wild-type and mutant Ant1 cause similar levels of cell death, as determined by flow cytometry after Annexin V-FITC and propoidium-iodide (PI) staining. Double negative cells are "non-apoptotic", Annexin V-positive PI-negative cells are "early apoptotic" and double positive cells are "dead". 24 hour treatment with 1 $\mu$M Staurosporine (STS) was used as a positive control. Data represented as mean +/- SEM.

**Fig. S7.**

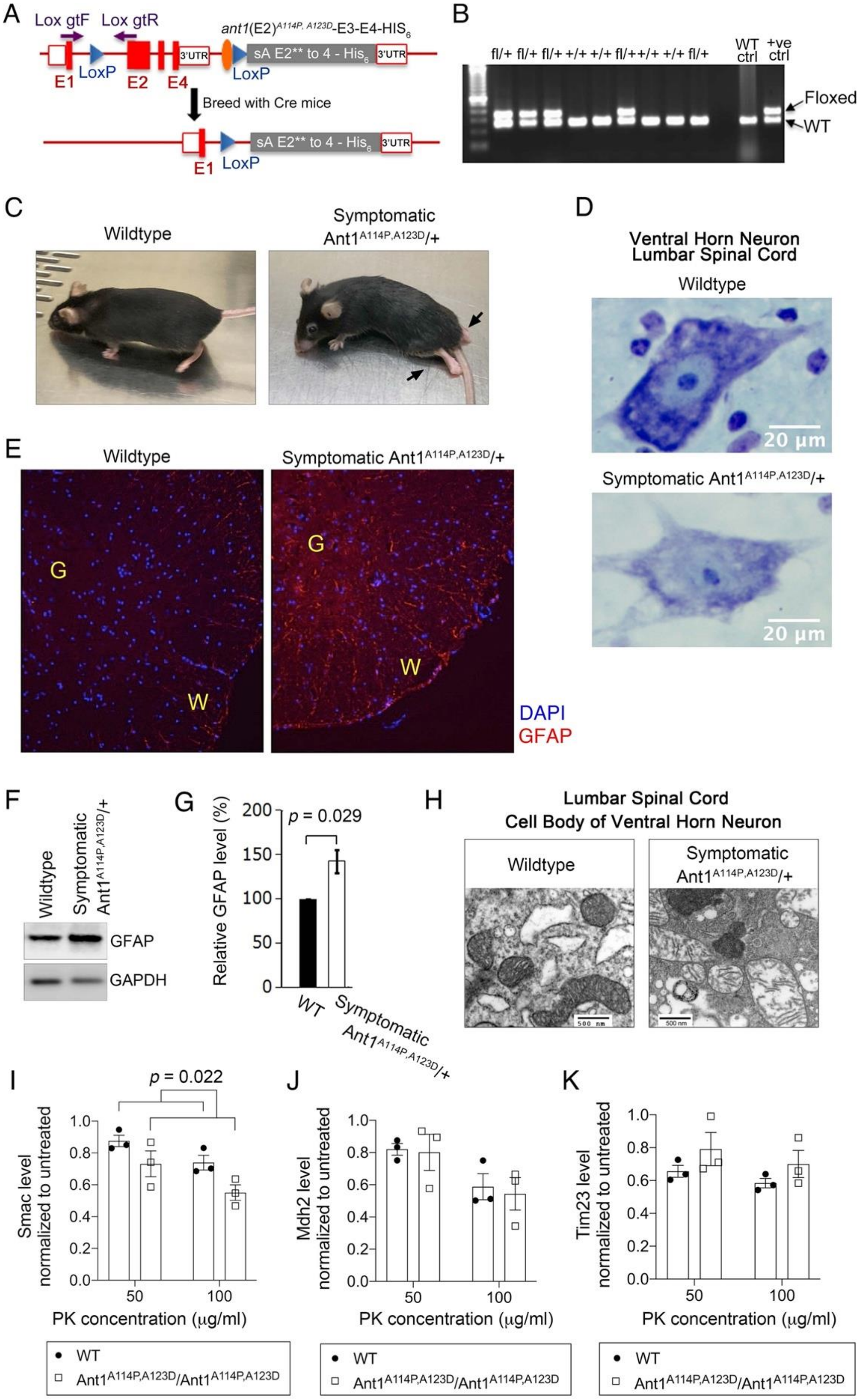
A
Lox gtF
Lox gtR
ant1(E2)A114P, A123D-E3-E4-HIS6
3'UTR
sA E2** to 4 - His6
LoxP
E1
E2
E4
Breed with Cre mice
B
fl/+ fl/+ fl/+ +/+ +/+ fl/+ +/+ +/+ fl/+
WT ctrl
+ve ctrl
Floxed
WT
C
Wildtype
Symptomatic Ant1A114P,A123D/+
D
Ventral Horn Neuron
Lumbar Spinal Cord
Wildtype
20 μm
Symptomatic Ant1A114P,A123D/+
E
Wildtype
Symptomatic Ant1A114P,A123D/+
G
W
DAPI
GFAP
F
Wildtype
Symptomatic Ant1A114P,A123D/+
GFAP
GAPDH
G
p = 0.029
Relative GFAP level (%)
0
50
100
150
200
WT
Symptomatic Ant1A114P,A123D/+
H
Lumbar Spinal Cord
Cell Body of Ventral Horn Neuron
Wildtype
Symptomatic Ant1A114P,A123D/+
500 nm
I
p = 0.022
Smac level normalized to untreated
PK concentration (μg/ml)
50
100
J
Mdh2 level normalized to untreated
K
Tim23 level normalized to untreated
0.0
0.2
0.4
0.6
0.8
1.0
WT
Ant1A114P,A123D/Ant1A114P,A123D

**Fig. S7. Ant1$^{A114P,A123D}$ knock-in mouse generation, neurodegeneration, and protease protection of homozygous Ant1$^{A114P,A123D}$/Ant1$^{A114P,A123D}$ mitochondria. (A)** Schematic of the strategy by which the knock-in *Ant1$^{A114P,A123D}$* mice were generated. E1 – E4, exons 1—4 of *ANT1*. In gray is the inserted cDNA containing two missense mutations in exon 2, followed by the endogenous 3' UTR. Lox gtF and gtR indicate genotyping primers. **(B)** Agarose gel electrophoresis of PCR genotyping using genotyping primers indicated in (A). Fl, floxed. **(C)** Ascending paralytic phenotype of an *Ant1$^{A114P,A123D}$*/+ mouse at 11-month old of age, and its wild-type littermate. Arrows point to paralyzed hindlimbs. **(D)** Nissl-stained lumbar spinal cord neuron of a symptomatic *Ant1$^{A114P,A123D}$*/+ mouse and wild-type littermate. This neuron shows loss of Nissl substance and blurring of nuclear boundaries, process known as "chromatolysis", which indicates neuron degeneration. **(E)** Indirect immunofluorescence for the astrocyte marker glial fibrillary acidic protein (GFAP) indicating spinal cord gliosis in a symptomatic *Ant1$^{A114P,A123D}$*/+ mouse. G, gray matter; W, white matter. **(F)** Immunoblot analysis of spinal cord lysate confirmed increase in GFAP in symptomatic *Ant1$^{A114P,A123D}$*/+ mice. **(G)** Quantitation from (F) showing significant increase in GFAP in the spinal cord of a symptomatic *Ant1$^{A114P,A123D}$*/+ mouse indicating neuroinflammation. *P* value was calculated from Student's *t* test. **(H)** Transmission electron microscopy of a ventral horn neuron of a symptomatic *Ant1$^{A114P,A123D}$*/+ mouse and wild-type littermate control. **(I – K)** Quantitation of immunoblot analysis of protease protection assay in wild-type and homozygous *Ant1$^{A114P,A123D}$*/ *Ant1$^{A114P,A123D}$* muscle mitochondria, as shown in Figure 7D. N = 3 mice per genotype. *P* values of genotype as main effect were calculated using a 2-way ANOVA. PK, proteinase K. Data represented as mean +/- SEM.

**Movie S1.**

Paralytic phenotype of an Ant1$^{A114P,A123D}$/+ mouse at the age of 12 months.

**Movie S2.**

Paralytic phenotype of an Ant1$^{A114P,A123D}$/+ mouse at the age of 16 months.

**Data S1. (separate file)**

Proteins that preferentially interact with Aac2$^{A128P}$-HIS6, compare with Aac2-HIS6 in low salt conditions.

**Data S2. (separate file)**

Proteomic comparison of the cytosolic fraction of Ant1$^{A114P,A123D}$- versus Ant1-transfected HeLa cells.

**Data S3. (separate file)**

Proteomic comparison of the cytosolic fraction of skeletal muscle from Ant1$^{A114P,A123D}$/+ versus wildtype mice.
